# Computational Analysis of Human Cysteine Redox Proteoforms Reveals Novel Insights

**DOI:** 10.1101/2024.09.18.613618

**Authors:** James N. Cobley, Panagiotis N. Chatzinikolaou, Cameron Schmidt

## Abstract

Since cysteine redox proteoforms (*i*) are virtually unstudied, we derived novel insights by computationally analysing the human proteome. Our analysis revealed a vast, effectively infinite, theoretical *i* space housing 3.02 x 10^169^ unique cysteine redox proteoforms. For >80% and 99% of the human proteome, the *i* space comprises 6.83 x 10^8^ and 1.76 x 10^31^ unique proteoforms, respectively. The heterogenous distribution of the *i* space by gene ontology terms, suggests, but does not prove, functional speciation. To theoretically limit the number of cysteine redox proteoforms that can be “downloaded” from the abstract *i* “cloud”, we implement novel equations. Protein copy numbers limit the *i* space by 161-logs to 4.04 x 10^7^ unique cysteine redox proteoforms per HeLa cell. An immutable law: the number of cysteine redox proteoform molecules (*Ni*) must equal the number of cysteine-containing protein molecules. We compute an *Ni* value of 1.70 x 10^9^ per HeLa cell. While *Ni* will be displaced from thermodynamic equilibrium towards the reduced state (e.g., ≈90%-reduced), it is possible that the number of partially oxidised cysteine redox proteoform molecules is in the order of 10^6-8^ per HeLa cell. Consistent with this, 100%-oxidised forms were observed in 60% of the proteins studied to date. Our analysis advances understanding of redox biology at the proteoform level.

## 1. Introduction

Cysteine redox proteoforms (*i*) include all the cysteine residue-and redox phase space-defined forms a protein molecule can adopt per equation 1 [1]. On an abstract mathematical plane, a protein molecule can be conceived as a superimposition of all possible *i* states [2–4], each with its own probabilities of being observed in a given biological system. For example, a monothiol molecule can be conceived as being in a superimposition of the reduced (0) and oxidised (1) states in binary redox phase space (i.e., [0 & 1]), where the <u>oxi</u>dised <u>form</u> is an oxiform per equation 2.

**Equation 1:** *i* = *n*^R^ where *n* and *R* are the cysteine redox phase space and residue values, respectively. Equation 1 can be automated to calculate an *i* value for a [Uniprot](https://www.uniprot.org/) protein accession of interest using our Redox_proteoforms_shinyapp/ shiny app.

**Equation 2:** *Oxiforms* = *i* − 1.

While all the theoretical *i* states are possible over time, a monothiol molecule cannot be reduced and oxidised simultaneously. At the single-molecule level, we can observe [0] or [1], but not [0 & 1]. For a molecule with 2 cysteines, we could observe [0,0], [0,1], [1,0] or [1,1], but not their superimposition ([0,0;0,1;1,0;1,1]). Importantly, these observable *i* states, the proteoforms that exist in a given biological system, govern:

1. **Cysteine biology**. Cysteine-dependent functions, such as transition metal ion binding, require specific redox states [5–7]. They can be reduced states, such as deprotonated thiolates (-2) for nucleophilic catalysis in the protein tyrosine phosphatase [PTP1B](https://www.uniprot.org/uniprotkb/P18031/entry), or oxidised states, such as the thiyl radical (+1) in [ribonucleotide reductase](https://www.uniprot.org/uniprotkb/P23921/entry).
2. **Redox regulation**. Post-translational control of proteins involves specific cysteine redox state transitions [8–12]. For example, PTP1B is inactivated when its active site cysteine attacks an oxygen atom in hydrogen peroxide (H_2_O_2_) [13,14], a reactive oxygen species (ROS) [15–17]. The sulfenic acid (RSOH) so-formed cannot dephosphorylate tyrosine.
3. **Oxidative stress**. The redox state of the cysteine proteome is determined by cysteine redox proteoforms. Hence, the cysteine proteome states associated with (causally or otherwise) oxidative stress, be it eustress or distress [18–20], are set by cysteine redox proteoforms [21].

Despite their importance, these fundamental protein units of redox biology are virtually unstudied [22]. Hence, we computationally studied cysteine redox proteoforms by quantifying the (a) theoretical number of unique cysteine redox proteoforms (the *i* space), (b) number of unique cysteine redox proteoforms that can form in a cell, and (c) number of cysteine redox proteoforms that do form. Our analysis advances understanding of redox biology at the proteoform level.

## 2. Results and discussion

### 2.1. Quantifying the number of theoretical human cysteine redox proteoforms

Python scripts calculated that the reference human proteome contained 262,025 cysteine residues spanning 19,817 proteins, representing 97.0% of the total 20,434 proteins [23–26]. To calculate the maximum number of theoretical human cysteine redox proteoforms, we used a script to sort human proteins by their number of cysteine residues. The output quantified the number of proteins with “*n*” cysteine residues (supplementary data file 1). For example, 888 proteins possessed 1 cysteine residue. After solving equation 1 for each cysteine residue class (e.g., 2 cysteines: *i* = 4), we multiplied the resulting *i* value by the number of proteins in the class (e.g., 2 cysteine residue class = *i*-4 x 1,140proteins). We then summed these *i* values to compute the maximum number of theoretical human cysteine redox proteoforms per equation 3.

**Equation 3.** ∑*i* ∗ *c* where *i* is the theoretical number of cysteine redox proteoforms that are possible for the number of cysteines in the class. Each *i* value is multiplied by the number of proteins in the cysteine residue class (*c*). The resulting cysteine residue class *i* values are summed to compute the theoretical limit.

Equation 3 set a theoretical limit of 3.02 x 10^169^ human cysteine redox proteoforms in binary redox phase space, a number so vast as to be effectively infinite. For context, if one counted a proteoform per second, then it would take 9.57 x 10^161^ years to count all of the theoretically possible human cysteine redox proteoforms, vastly exceeding the estimated age of the universe ≈1.38 x 1010 years. For reference, many other species possess vast i spaces (Figure 1, Table 1, & supplementary data file 2).

**Figure 1.**
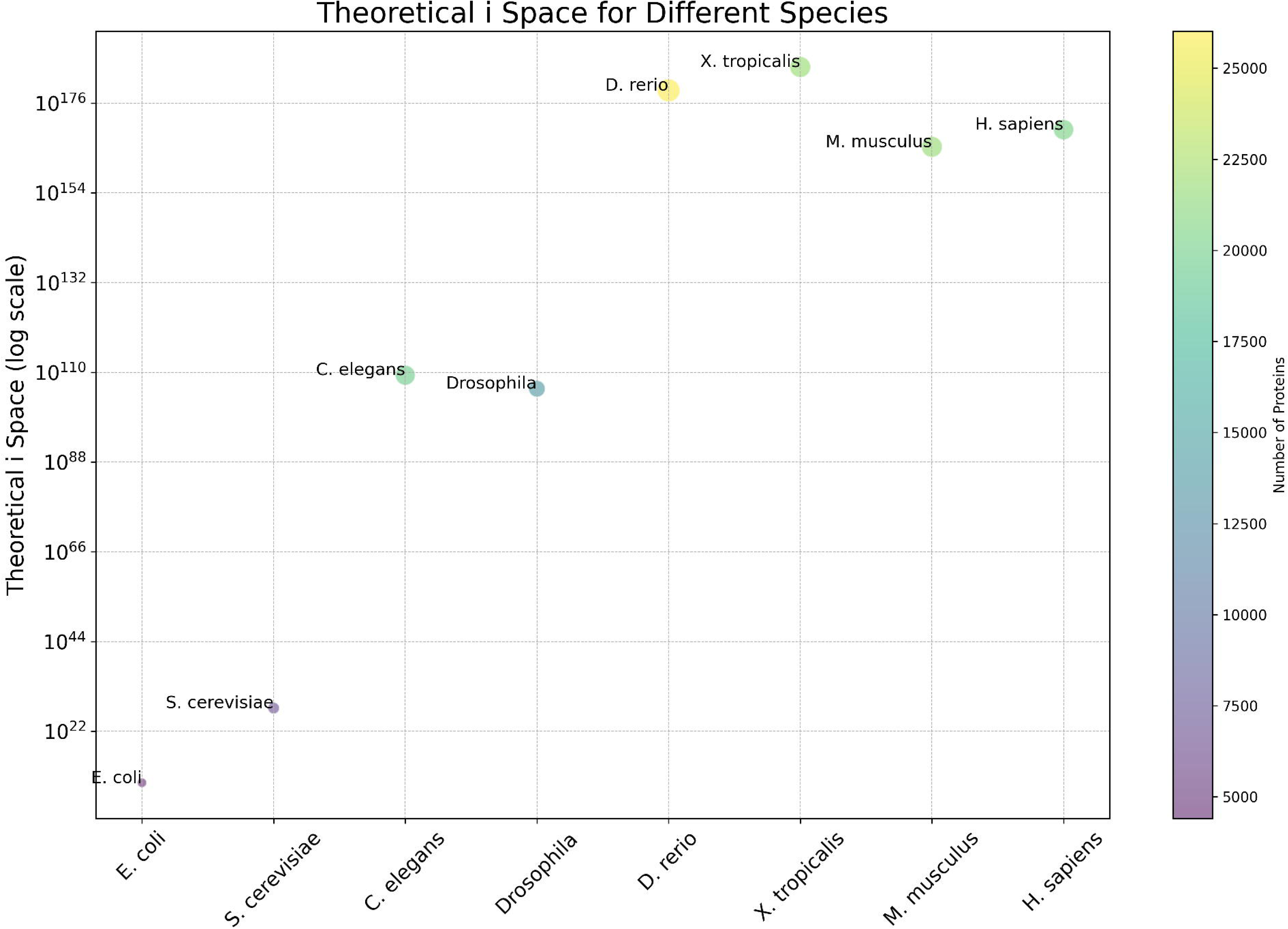
Theoretical *i* Space for Different Species. The bubble plot illustrates the theoretical *i* space across various species. The y-axis represents the theoretical *i* space on a log scale, while the x-axis lists the species. Bubble sizes correspond to the number of proteins in each species, and the colour gradient reflects the same. To fit the bubbles on the graph two extreme outliers were omitted: M9PB30 with 2,647 cysteine residues in *Drosophila* and A0A8M9P2Z8 with 1,124 cysteine residues in *D. rerio*.

**Table 1.**
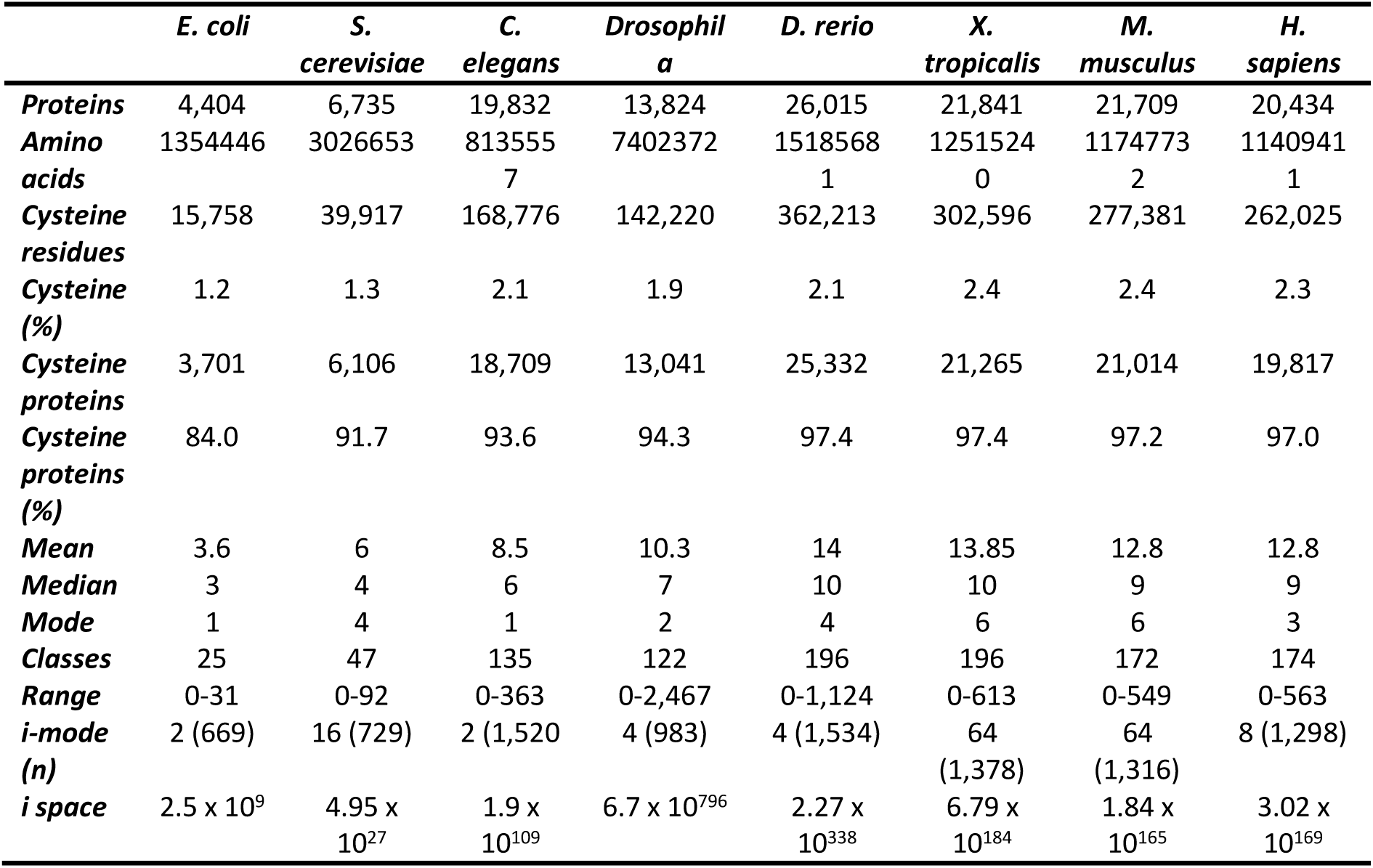
Quantitative information on the selected proteome parameters, such as the number of proteins, by selected model species. Cysteine residue mean, median, and mode values are displayed. Data were extracted by analysing their reference proteomes per the methods section.

In sum, equation 3 defined an upper limit of 3.02 x 10^169^ unique cysteine redox proteoforms in humans. This number is so large that it can for all practical purposes be treated as effectively infinite, especially when combined with all the other possible sources of protein speciation [27]. These other forms of speciation mean that even when the protein-specific *i*-space is finite, the number of possible combinations, such as those involving over 300 post-translational modifications across hundreds of amino acids, is still effectively infinite.

### 2.2. Defining the composition of the human *i* space

Next, we defined the composition of the abstract human *i* space. Since equation 3 summed class-specific *i* values, we defined the relationship between the cysteine residue and protein integers in the human proteome. The number of cysteine residues spanned 2-logs from 0 to 563 residues. Consistent with a heterogenous protein frequency pattern, we observed a marked leftward shift in the distribution of cysteine residues (Figure 2A, Table 2, & supplementary data file 1). Most proteins possessed <20 cysteine residues. The distribution exhibited a long tail, reflecting the presence of relatively rare proteins with >100 cysteine residues. Ergo, the share of each cysteine residue class to the overall percentage of proteins in the proteome decreased as an inverse function of the cysteine residue count (Figure 2B).

**Figure 2.**
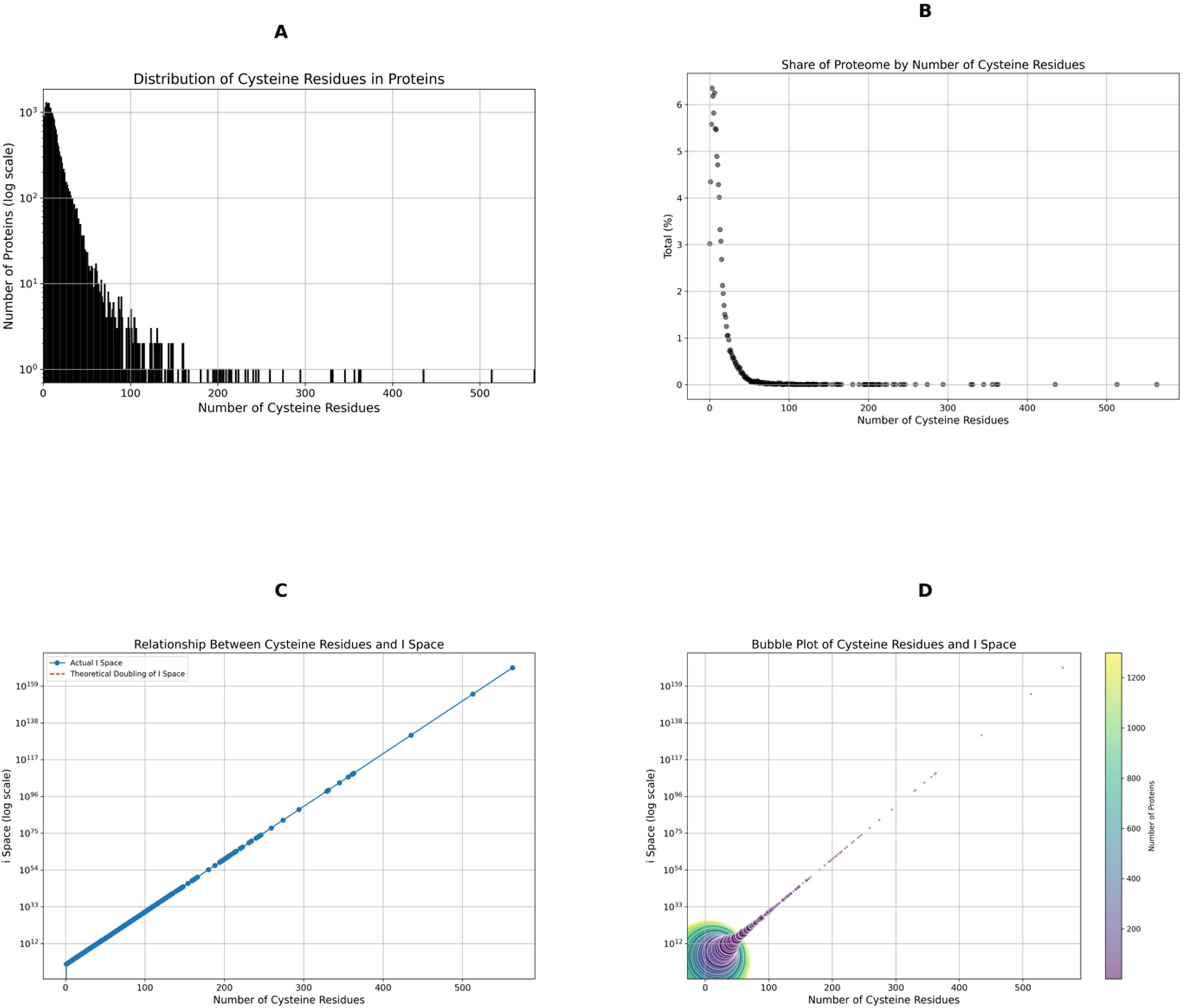
Distribution and Theoretical *i* Space Analysis of Cysteine Residues in the Human Proteome. (A) Histogram displaying the distribution of cysteine residues across proteins in the reference human proteome, revealing a marked leftward shift, with most proteins containing fewer than 20 cysteine residues. The long tail reflects relatively rare proteins with more than 100 cysteine residues. (B) Share of Proteome by Number of Cysteine Residues illustrating the inverse relationship between the number of cysteine residues and the percentage share of the proteome. Proteins with a high number of cysteine residues have a lower percentage share of the proteome, while those with fewer cysteine residues account for a larger share. (C) Relationship Between Cysteine Residues and *i* Space. The plot demonstrates the theoretical doubling of the *i* space with increasing cysteine residues. The linear relationship on a log scale confirms that the *i* space doubles with each additional cysteine residue. (D) Bubble Plot of Cysteine Residues and *i* Space. This bubble plot visualizes the relationship between cysteine residues and the *i* space, with the bubble size representing the number of proteins. The x-axis shows the number of cysteine residues, and the y-axis (log scale) indicates the *i* space. The colour gradient, from purple to yellow, corresponds to the number of proteins, highlighting that a small number of proteins with high cysteine counts contribute disproportionately to the total *i* space.

**Table 2.** Reference values by cysteine residue classifier range in the human proteome. The reference values cover the number of proteins and cysteine residues in each range, their percentages of the overall total, and the *i* value for each range.

| Classifier range | Proteins | Proteins (%) | Cysteine residues | Cysteine residues (%) | *i* |
| --- | --- | --- | --- | --- | --- |
| 1-10 | 11254 | 61113 | 61113 | 2.08E+06 | 2.08E+06 |
| 11-20 | 5334 | 77040 | 77040 | 6.81E+08 | 6.81E+08 |
| 21-30 | 1683 | 41657 | 41657 | 2.64E+11 | 2.64E+11 |
| 31-40 | 784 | 27388 | 27388 | 1.32E+14 | 1.32E+14 |
| 41-50 | 328 | 14670 | 14670 | 5.33E+16 | 5.33E+16 |
| 51-60 | 135 | 7494 | 7494 | 3.42E+19 | 3.42E+19 |
| 61-70 | 84 | 5472 | 5472 | 1.89E+22 | 1.89E+22 |
| 71-80 | 48 | 3633 | 3633 | 1.26E+25 | 1.26E+25 |
| 81-90 | 41 | 3534 | 3534 | 1.20E+28 | 1.20E+28 |
| 91-100 | 21 | 2036 | 2036 | 9.66E+30 | 9.66E+30 |
| 101-120 | 29 | 3129 | 3129 | 1.69E+36 | 1.69E+36 |
| 121-140 | 26 | 3338 | 3338 | 8.44E+41 | 8.44E+41 |
| 141-160 | 13 | 1953 | 1953 | 4.41E+48 | 4.41E+48 |
| 161-180 | 4 | 671 | 671 | 1.53E+54 | 1.53E+54 |
| 181-200 | 6 | 1171 | 1171 | 2.18E+60 | 2.18E+60 |
| 201-250 | 15 | 3303 | 3303 | 1.29E+74 | 1.29E+74 |
| 251-300 | 3 | 827 | 827 | 3.18E+88 | 3.18E+88 |
| 301-400 | 6 | 2085 | 2085 | 2.36E+109 | 2.36E+109 |
| 401-500 | 1 | 435 | 435 | 8.87E+130 | 8.87E+130 |
| 501-600 | 2 | 1076 | 1076 | 3.02E+169 | 3.02E+169 |

The implications of the observed cysteine distribution pattern on the composition of the abstract *i* space are profound as the *i* space doubled each time the cysteine residue integer increased by 1. For example, the *I* space doubled from 2 to 4 unique cysteine residues for proteins with 1 and 2 cysteine residues, respectively (supplementary data file 1). Hence, we observed a perfect linear relationship between the size of the *i* space and the protein cysteine residue integer (Figure 2C). This linear relationship revealed a disproportionate influence of just a few proteins on the overall composition of the *i* space (Figure 2D). One protein accounted for essentially the entire 99.9% recurring *i* space: With 563 cysteine residues, [Spondin](https://www.uniprot.org/uniprotkb/A2VEC9/entry) can form 3.02 x 10^169^ cysteine redox proteoforms. This number is so large that the notational value stayed at 3.02 x 10^169^ when we added all the other *i* values to it because of the 15-log difference between the *i* values for spondin and [titin](https://www.uniprot.org/uniprotkb/Q8WZ42/entry) (Cys-513). One must subtract the 3 proteins with the most cysteine residues (spondin, titin, & [IgGFc-binding protein](https://www.uniprot.org/uniprotkb/Q9Y6R7/entry), Cys-435) before the summed total for the remaining 19,813 proteins changes from the notational one for [fibrillin-2](https://www.uniprot.org/uniprotkb/P35556/entry) (Cys-363):

**The *i* space for 19,813 proteins = 2.36 x 10^109^**. [Fibrillin-2 (1.88 x 10^109^) + all other class-specific *i* values]

To provide reference values, we stratified the cysteine residue distribution ranges using 10, 20, 50, and 100 as the cut-off classifiers for 1-100, 101-200, 200-300, and >301 cysteine residues, respectively (Table 2). Most of the human cysteine proteome (57.8%) contained between 1-10 cysteine residues (23.3% of the total). The 26.9% of proteins with 11-20 cysteine residues accounted for 29.4% of all the cysteine residues. Combined the 1-20 cysteine residue ranges represented 83.7% and 52.7% of the proteins and cysteine residues in the human cysteine proteome, respectively. The theoretical *i* space for >80% of the human proteome is 6.83 x 10^8^. Only 105 proteins (0.052% of the proteome) possessed >100 cysteine residues. Of these, 27 proteins had >200 cysteine residues. As a result, the theoretical *i* space for 99.95% of the human proteome is 1.76 x 10^31^.

In sum, the mathematics of equation 1 unmasked the disproportionate influence of a small number of proteins with large cysteine residue integers on the composition of the human *i* space. Hence, the maximum *i* space for >80% of the human proteome is a striking 161-logs smaller than the total *i* space: 6.83 x 10^8^ vs. 3.02 x 10^169^ cysteine redox proteoforms.

### 2.3. Defining the distribution of the human *i* space using Gene Ontology terms

To define the distribution of the *i* space by Gene Ontology (GO) terms, we listed Uniprot accessions for the proteins in each cysteine residue class (e.g., the 1 cysteine column lists 888 accessions) using a custom script (supplementary data file 3). Our script matched the accessions to selected GO terms for cellular location by parsing the file and the UniProt database. The output (supplementary data file 4) displayed the number of proteins in each term and sorted them into cysteine residues classes, which enabled the location-specific *i* space values to be calculated per equation 3. The extracellular matrix harboured the largest *i* space (3.02 x 10^169^) due to the presence of spondin (Figure 3A). While the extracellular milieu is an important redox hub [28,29], comparatively little is known about its cysteine redox biology. For example, the function of spondin remains mysterious. Its role in neuronal aggregation [30] may involve its 103 disulfide bonds (Prosite rule) and/or the 357 “free” cysteines. Are they a sponge for soaking up oxidative stress in the brain? [31] Or a store of oxidising equivalents for relaying redox signals?

**Figure 3.**
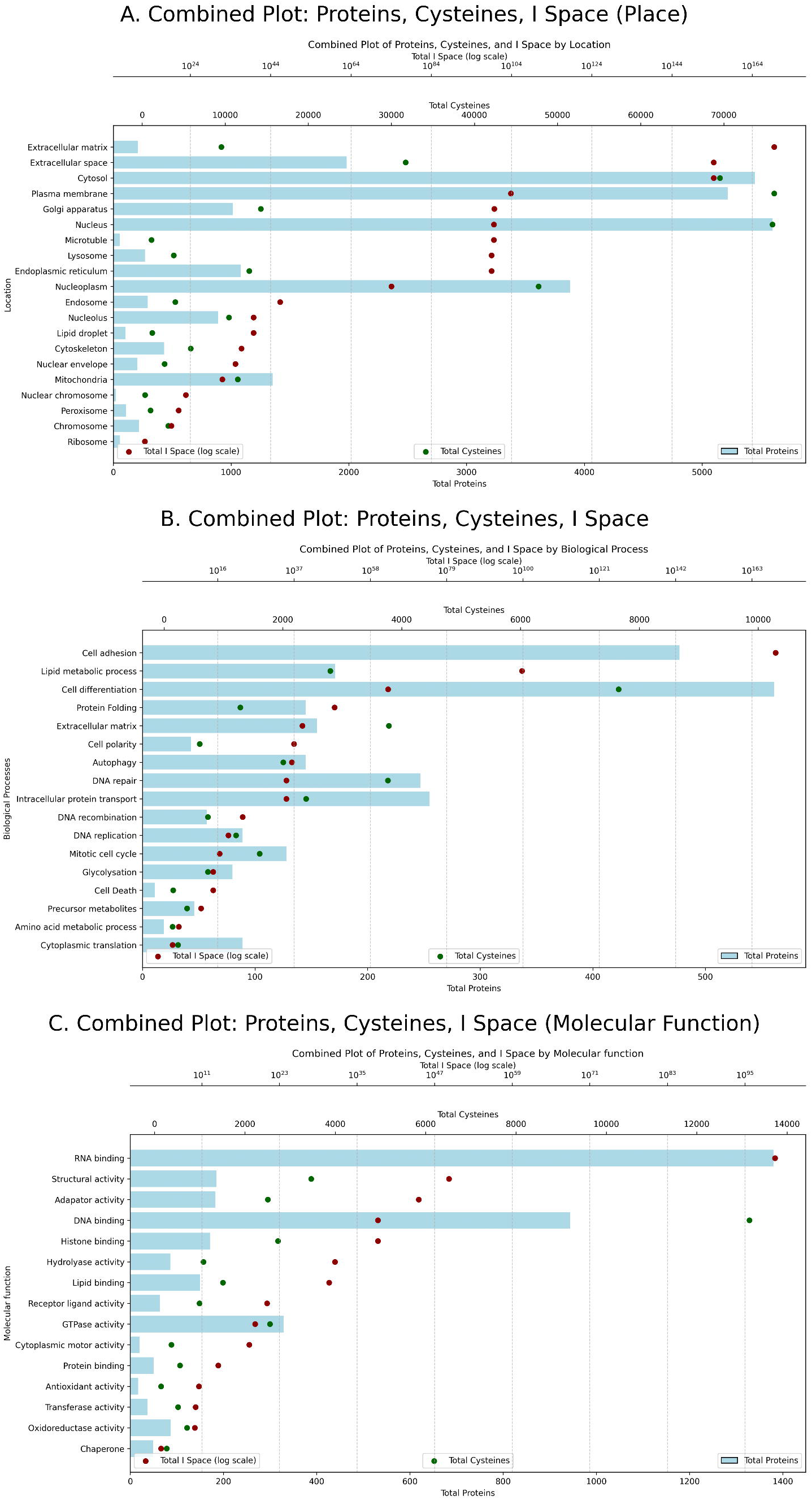
Plotting the *i* space by GO terms. Panels A, B, and C displays the distribution of total proteins, total cysteines, and theoretical *i* space for selected GO terms corresponding to location, biological process, and molecular function, respectively. In each panel, the blue bars, green dots, and red dots represent total proteins, total cysteines, and the theoretical *i* space (log scale) respectively.

Within the cell, the cytosol housed the largest theoretical *i* space at 2.68 x 10^154^. The location-specific *i* space values for the plasma membrane (7.2 x 10^103^) and nucleus (4.4 x 10^99^) were also appreciable. Hence, redox regulation hubs can theoretically house multitudes of diverse cysteine redox proteoforms. Conversely, the ribosome housed the smallest theoretical *i* space of 4.4 x 10^12^. The intracellular location-specific *i* space ranges spanned 146-logs. While these differences exist in the abstract mathematical plane, the relative volume of each location is likely to determine their biologically accessible *i* space. For example, at an equal protein density, the cytosol can house more proteoforms compared to the ribosome due to a 560,000 times volume differential (cytosol ≈ 1,400 μm^3^ vs. ribosome ≈ 2.5 × 10^−3^ μm^3^).

Next, we plotted the abstract *i* space by selected GO terms for biological processes (Figure 3B, supplementary data file 5). Cell adhesion possessed the largest process-specific *i* space of 3.02 x 10^169^ due to the presence of spondin. Additionally, 12 (11.4%) of proteins with >100 cysteine residues are involved in cell adhesion processes. Cysteine-rich proteins seem to be important molecular glues, potentially due to disulfide bonds. The process-specific *i* space values associated with lipid metabolic (5.5 x 10^99^) and cell differentiation (8.23 x 10^62^) were considerable. This suggests the intuitively appealing idea of a role for cysteine redox proteoform speciation in cell differentiation. Conversely, many biological processes had *i* values <10^23^. For example, the process-specific *i* value for cytoplasmic translation was 4,220. Many tightly controlled processes, such as DNA recombination, had comparatively compact abstract *i* spaces. While speculative, there may be equally tight chemical strictures placed on cysteine redox proteoform speciation when molecular fidelity is paramount.

Finally, we plotted the abstract *i* space by selected GO terms for molecular function (Figure 3C, supplementary data file 6). In sharp contrast to the ribosome and cytoplasmic translation, RNA binding (4.4 x 10^99^) possessed the largest function-specific *i* value. For example, the RNA binding protein [NBPFA](https://www.uniprot.org/uniprotkb/Q6P3W6/entry) contains 139 cysteine residues (*i* = 6.97 x 10^41^). It would be fascinating to determine whether these cysteines regulate RNA binding. In line with the importance of disulfide bonds, structural activity (1.75 x 10^49^) possessed the next largest function-specific *i* value. For example, 106 (83.4%) of the 127 cysteines in [LAMB1](https://www.uniprot.org/uniprotkb/P07942/entry) form disulfide bonds. Conversely, chaperone activity (49,154) possessed the smallest function-specific *i* value. We speculate that a more compact *i* space benefits molecular fidelity by limiting the potential for protein speciation.

In sum, our analysis is congruent with Figure 2 and Table 2. Equation 1 produced large discrepancies between the number of proteins and cysteines in a GO term and the *i* space. For example, spondin alone is all that is required for a GO term to house the largest *i* space in humans. Supplementary data files 3-6 define a useful resource for follow-up computational and experimental analyses. For example, investigations to determine the relationship between the size of the abstract *i* space and the number of biologically accessed cysteine redox proteoforms. The heterogeneous distribution of the *i* space suggests, but does not prove, functional speciation.

### 2.4. Mathematically limiting the biologically accessible *i* space

To define the factors that limit the biologically accessible *i* space, we developed novel equations. The presence of specific proteins may dictate the theoretical *i* space in a given biological system, such as a tissue. For example, the lack of [p53](https://www.uniprot.org/uniprotkb/P04637/entry) would decrease the theoretical *i* space by 1,024 proteoforms on expression grounds alone. Hence, the theoretical limit for a given biological system partially depends on the identity of the proteins that are expressed per equation 4.

**Equation 4**: ∑*i* ∗ *cx* where *i* is the number of cysteine redox proteoforms times the number of proteins in the cysteine residue distribution class that are expressed in the biological system (*cx*). The *cx* values are then summed to calculate the theoretical *i* space limit for a given biological system.

To determine whether equation 4 limited the theoretical *i* space, we analysed a proteomic HeLa cell dataset [32] using a script that matched proteins to Uniprot accessions. The 9,906 matched proteins (supplementary data file 7) and their 128,541 cysteine residues represent 48.5% and 49.1% of the human proteome and cysteine proteome, respectively. Less cysteine residues classes (-38 classes) and proteins (-50%) concomitantly limited the *i* space per class vs. the reference human proteome. For example, the theoretical *i* space limit for the 1-20 residue range contracted by 50.6% from 6.84 x 10^8^ in the reference proteome to 3.46 x 10^8^ in HeLa cells (Table 3). Equation 4 constrained the theoretical *i* space by 15-logs from 3.02 x 10^169^ to 2.68 x 10^154^ in Hela cells vs. the reference human proteome due to the absence of spondin.

**Table 3.** The theoretical *i* space limit for each cysteine residue class between 1-20 in the reference human proteome and a published HeLa cell proteome. Less proteins in the HeLa proteome vs. the reference decrease the *i* space in each class on expression grounds per equation 4.

| Cysteine residue class | Reference | | HeLa | | Hela vs. ref $i$ space (% decrease) |
| --- | --- | --- | --- | --- | --- |
| | Proteins | $i$ space | Proteins | $i$ space | |
| 1 | 888 | 1.78E+03 | 504 | 1.01E+03 | -56.76 |
| 2 | 1140 | 4.56E+03 | 566 | 2.26E+03 | -49.65 |
| 3 | 1298 | 1.04E+04 | 606 | 4.85E+03 | -46.69 |
| 4 | 1263 | 2.02E+04 | 607 | 9.71E+03 | -48.06 |
| 5 | 1189 | 3.80E+04 | 567 | 1.81E+04 | -47.69 |
| 6 | 1278 | 8.18E+04 | 618 | 3.96E+04 | -48.36 |
| 7 | 1121 | 1.43E+05 | 532 | 6.81E+04 | -47.46 |
| 8 | 1116 | 2.86E+05 | 578 | 1.48E+05 | -51.79 |
| 9 | 999 | 5.11E+05 | 446 | 2.28E+05 | -44.64 |
| 10 | 962 | 9.85E+05 | 404 | 4.14E+05 | -42.00 |
| 11 | 876 | 1.79E+06 | 378 | 7.74E+05 | -43.15 |
| 12 | 820 | 3.36E+06 | 361 | 1.48E+06 | -44.02 |
| 13 | 679 | 5.56E+06 | 317 | 2.60E+06 | -46.69 |
| 14 | 628 | 1.03E+07 | 270 | 4.42E+06 | -42.99 |
| 15 | 548 | 1.80E+07 | 233 | 7.63E+06 | -42.52 |
| 16 | 434 | 2.84E+07 | 207 | 1.36E+07 | -47.70 |
| 17 | 399 | 5.23E+07 | 207 | 2.71E+07 | -51.88 |
| 18 | 347 | 9.10E+07 | 164 | 4.30E+07 | -47.26 |
| 19 | 308 | 1.61E+08 | 154 | 8.07E+07 | -50.00 |
| 20 | 295 | 3.09E+08 | 156 | 1.64E+08 | -52.88 |

When it is expressed, protein concentration may dictate the biologically accessible *i* space. For example, even an unrealistically high concentration of 1 mole of spondin, would contract the theoretical *i* space by 146-logs from 3.02 x 10^169^ to 6.022 x 10^23^. More realistic spondin concentrations of 1 micromole, 1 picomole, 1 femtomole, and 1 attomole would collapse the biologically accessible *i* space to 6.022 x 10^17^, 6.022 x 10^11^, 6.022 x 10^8^, and 6.022 x 10^5^ cysteine redox proteoforms, respectively. The latter value represents a 164-log reduction from the theoretical maximum, reflecting the profound importance of protein concentration values, as codified in equation 5.

**Equation 5.** ∑_*index*_ (*min*(*i*, *N*)) where *i* is the protein cysteine redox proteoform integer *N* is the protein copy number. The minimum values are summed to calculate the maximum number of biologically accessible cysteine redox proteoforms.

To determine whether equation 5 limited the theoretical *i* space, we retrieved a cysteine residue number from the reference human proteome for the 8,099 proteins with estimated protein copy numbers (*N*) per HeLa cell [32] (supplementary data file 8). The summed *N* value totalled 2.05 x 10^9^ protein molecules per HeLa cell. Compared to equation 4, equation 5 constrained the theoretical *i* space by 147-logs from a maximum of 2.68 x 10^154^ to 4.04 x 10^7^ unique cysteine redox proteoforms per HeLa cell. Hence, *N* is the dominant factor responsible for constraining the biologically accessible *i* space in a HeLa cell, a finding that is likely to be generally applicable to most systems.

Interestingly, the abstract *i* space available to the 10 most abundant proteins that represent 10.5% of all the protein molecules in a HeLa cell is just 64 unique cysteine redox proteoforms. Five proteins (50%) contained a cysteine residue (3 cysteines = 2, 4 cysteines = 3). Bar [TAGLN2](https://www.uniprot.org/uniprotkb/P37802/entry), each protein can be classed as an antioxidant ([PRDX1](https://www.uniprot.org/uniprotkb/Q06830/entry), [GSTP1](https://www.uniprot.org/uniprotkb/P09211/entry)) and/or redox regulated ([GAPDH](https://www.uniprot.org/uniprotkb/P04406/entry), [CFL1](https://www.uniprot.org/uniprotkb/P23528/entry)) molecule. As Cys124 is >20% oxidised in 5 different mouse tissues [33], we speculate that TAGLN2 is redox regulated. TAGLN2 is ideal for testing our proposal that a high *N* is associated with greater biologically accessed *i* space compared to a protein with a low *N* at a set cysteine residue integer.

Abundant proteins tended to have a lower theoretical *i* space (Figure 4A-B). For example, the 1,000 most abundant proteins accounted for 85% (1.70 x 10^9^) of the total protein molecules in the cell. However, their theoretical *i* space (2.88 x 10^17^) accounted for just 1.25217 x10^−90^% of 2.3×10^109^. Consistent with this, 94.5% (945) of these proteins contained <10 cysteine residues. Only 8 (0.8%) possessed >20 cysteine residues. Moreover, to the best of our knowledge, all of the proteins with a *k* >10^5^ M^-1^ s^-1^ with H_2_O_2_ were in the top 2,000 most abundant molecules. For example, all PRDX isoforms were in the top 1,000. Our observations suggested a design principle whereby the most abundant molecules contained <10 cysteine residues, implying that:

a. The percentage redox gradients associated with cysteine oxidation events are sharper. They range from 100 (1 cysteine) to 10% (10 cysteines).
b. These proteins are more likely to be oxidised than less abundant molecules, when all else is equal. The equality clause assumes an equivalence in all other important variables. When the clause is violated, the statement may no longer hold true.
c. Cysteine oxidation events are funnelled into a comparatively narrower set of possible proteoform coordinates. This funnel may help to encode specificity at the proteoform-level, channelling electrons into specific oxiforms.

**Figure 4.**
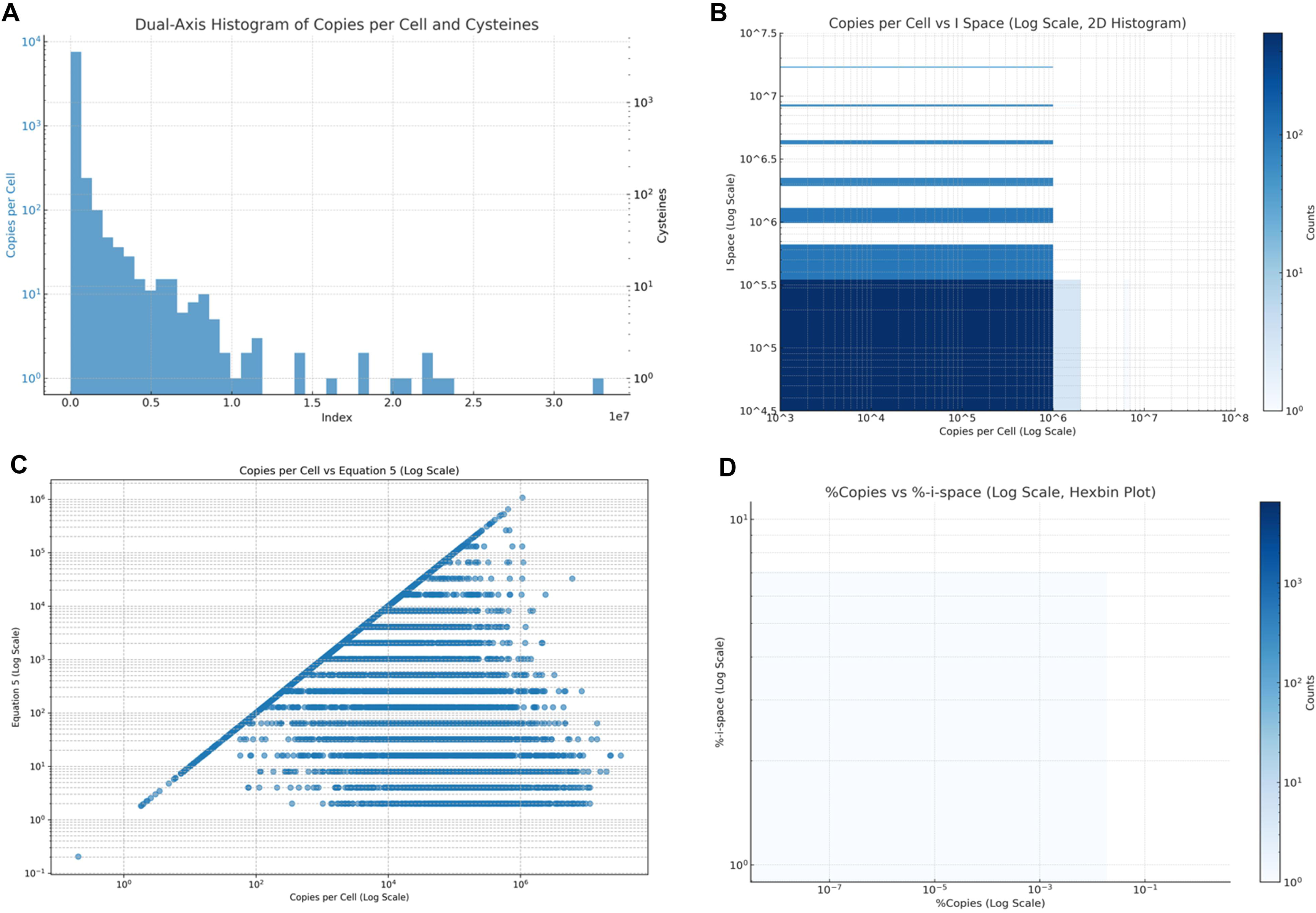
The biologically accessible *i* space in HeLa cells. A: *Dual-Axis Histogram of Copies per Cell and Cysteines*. This plot shows the distribution of protein copies per cell (left y-axis, blue bars) and the number of cysteines in proteins (right y-axis, black bars). The x-axis represents the index of the proteins. Both axes are on a linear scale. B: *Copies per Cell vs I Space*. This 2D histogram illustrates the relationship between the number of copies per cell and the I space of proteins. Both axes are on a logarithmic scale, with darker blue areas indicating higher counts of proteins in those bins. C: *Copies per Cell vs Equation 5.* This scatter plot displays the correlation between copies per cell and Equation 5 values for proteins, with both axes on a logarithmic scale. Each blue dot represents a protein, showing how these two metrics vary across different proteins. D: *%Copies vs %-i-space*. This hexbin plot depicts the relationship between the percentage of copies and the percentage of *i* space in proteins. Both axes are on a logarithmic scale. The colour intensity represents the density of data points, with darker blue indicating higher concentrations.

The theoretical *i* space was unconstrained by *N* for 5,399 (67%) proteins (Figure 4C). Conversely, *N* constrained the biologically accessible *i* space for 2,700 (33%) of proteins. The most profound constriction occurred for the 39 (0.48%) proteins with >100 cysteine residues. Equation 5 constrained their *i* space by 104-logs, from 2.3 x 10^109^ theoretical to 1.09 x 10^5^ biologically accessible unique cysteine redox proteoforms. Their impact was still outsized: 0.005% of the total protein molecules accounted for 0.27% of the biologically accessible *i* space. Despite representing 2,100 times more of the total protein molecules, the 10 most abundant proteins comprised just 0.0001% of the *i* space. Similarly, the 1,000 most abundant proteins (≈85% of the total molecules) accounted for just 16.8% of the biologically accessible *i* space (*i* = 6.8 x 10^6^). The top 10, 100, and 1,000 proteins by *i*-space share accounted for 12.3% (*i* = 5.07 x 10^6^), 40.3% (*i* = 1.63 x 10^7^), and 84.1% (*i* = 3.40 x 10^7^) of the total biologically accessible *i* space, respectively. None of these proteins contained <13 cysteine residue. Even though equation 5 does not constrain the space for most proteins, the biologically accessible *i* space is concentrated in 3.5% of the total protein molecules (Figure 4D).

In sum, most of the vast abstract *i* space cannot actually be biologically accessed because either the protein is not expressed or there are far too few molecules of an expressed protein to form all of the possible proteoforms. Novel equations limited the potential *i* space by 161-logs from a “cloud” value of 3.02 x 10^169^ to a “download limit” of 4.04 x 10^7^ unique cysteine redox proteoforms per Hela cell.

### 2.5. Factors controlling the number of cysteine redox proteoforms that are biologically accessed

Equation 4-5 limited the number of *unique* cysteine redox proteoforms in a given biological system. In every biological system, every protein molecule with a cysteine residue (*N_c_*) <u>is</u> a redox proteoform. Hence, the number of cysteine redox proteoforms (*N_i_*) must equal *N_c_*per equation 6. Equation 6 defined an *Ni* of 1.70 x 10^9^ cysteine redox proteoforms per HeLa cell, exceeding the number of unique forms by 2-logs. Formal equivalence means *N_c_* controls *N_i_*.

**Equation 6.** *N*_*i*_ = *N*_*c*_ where *N_i_* and *N*_c_ are the number of cysteine redox proteoforms and protein molecules with at least one cysteine residue, respectively. Ergo, *N_i_* cannot exceed *N_c_*.

What is *Ni* composed of? Each unique cysteine redox proteoform represents a potential microstate (Ω) of the system where each Ω can be related to entropy (*S*) via the Boltzmann equation (equation 7). Note, *S* is satisfied in both the forward (oxidising) and backward (reduction) directions. Each direction is intimately linked to energy-transduction pathways [34], such as the NADP^+^/NADPH redox couple [35]. There are many more ways for the Ω that make up the *N_i_* system to be partially vs. fully reduced or oxidised. Specifically, the number of fully vs. partially reduced or oxidised Ω in the human proteome differed by 165-logs: 3.96 x 10^4^ vs. 3.02 x 10^169^, respectively. Orders of magnitude more partial vs. fully reduced/oxidised Ω make it more probable to observe a protein in a partially reduced/oxidised proteoform state. Hence, probabilistic thermodynamics predicts that partially reduced/oxidised Ω dominate *Ni*.

**Equation 7.***S* = *k*_*B*_ *InΩ* where *k_B_* is the Boltzmann constant.

However, *N_i_* is likely to be displaced far from thermodynamic equilibrium in biological systems as manifested by the dominance of fully reduced or more reduced than oxidised Ω to *N_i_*. Indeed, *N_i_* is ≈90%-reduced in HeLa cells [36]. This bias towards the reduced state reflects two key factors. First, oxidation can only occur when electron transfer to the cysteine sulfur atom is possible. All reactants must be accessible. However, many cysteines are either partially or fully shielded from the solvent. For example, solvent accessible surface area (SASA) analysis of an AlphaFold [37,38] predicted structure of [PTEN](https://www.uniprot.org/uniprotkb/P23528/entry) revealed that 4 out of 10 cysteine residues were buried (Figure 5). If these residues remained inaccessible, 40% of the cysteines should be reduced. Decreasing the biologically accessible *i* space by 2-logs from 1,024 to 64 cysteine redox proteoforms. Hence, structural arguments can bias the biologically accessed *i* space toward the reduced state.

**Figure 5.**
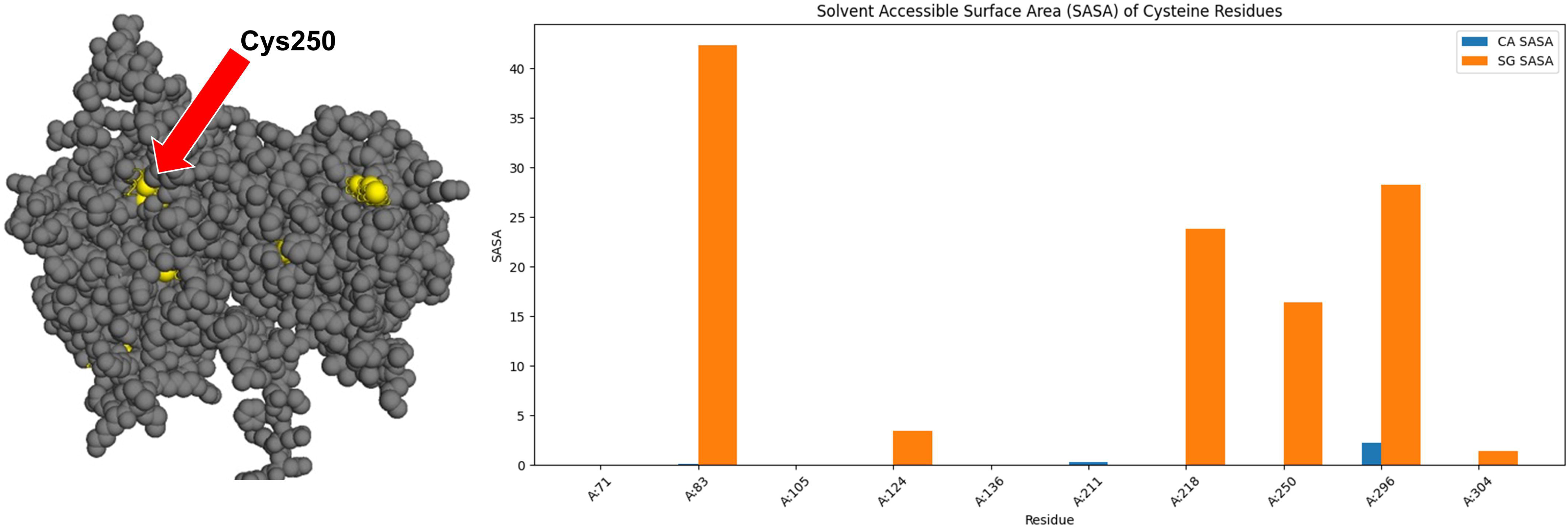
SASA analysis of PTEN. Left. A PyMOL screenshot of a PTEN structure with the cysteine residues highlighted in yellow. The block red arrow spotlights Cys250 as a partially exposed residue. In this structure 40% of the cysteines are buried. Right. Biopython calculated SASA values for the cysteine residues in PTEN.

Second, cysteine oxidation is under strict kinetic control [39]. While the phosphatase activity of PTEN can be inactivated when its active site cysteine residue (Cys124) attacks H_2_O_2_ to form a RSOH, the kinetic rate constant (*k*) is slow. Using published data [40], we estimated that a *k* of 5.717 M^-1^ s^-1^ at 25°C. With this *k* and 3188 PTEN molecules in a HeLa cell volume of 2.6 x 10^3^ μm^3^, a steady-state H_2_O_2_ concentration of 10 nM [41] would oxidise 1 PTEN molecule (0.03%) per hour per cell. However, a more precise computational approach using floating-point arithmetic revealed that, under these conditions, the oxidation rate is so low that effectively zero PTEN molecules would be oxidized over the same time period. Still, when Cys124-SOH does form, a fast condensation reaction (*k* ≈ 10^5^ M^-1^ s^-1^) with Cys71-S^-^ to form a disulfide bond would produce a 20%-oxidised cysteine redox proteoform (reactions 1 and 2). Assuming similar *k* values for the other cysteine residues, the formation of the 100%-oxidised PTEN-oxiform would seem quite unlikely at least in theory. Hence, kinetic arguments can bias the biologically accessed cysteine redox proteoforms toward the reduced state.

**Reaction 1**: Cys124-S^-^ + H_2_O_2_ Cys124-SOH + ^-^OH. Produces a 10%-oxidised PTEN proteoform.

**Reaction 2**: Cys124-SOH + Cys71-S^-^ Cys124-S-S-Cys71 + H^+^. Produces a 20%-oxidised PTEN proteoform.

When considering structure and kinetic-based arguments one should note that many partially exposed and even buried cysteine residues “sluggish” H_2_O_2_ reactivity can be oxidised [33,42,43]. These empirical observations point to the importance of dynamic conformational landscapes and a broad vista of cysteines oxidation mechanisms. As a result, even a highly reduced *N_i_* system can accommodate many Ω with at least one oxidised cysteine residue. For example, simply computing 10% of *Ni* revealed that there could be 1.7 x 10^8^ partially oxidised cysteine redox proteoform molecules per Hela cell. Conservatively, revising the number down by 2-logs, still gave 1.7 x 10^6^ cysteine redox proteoforms per Hela cell. For reference, the latter value is 6.5 times larger than the number of cysteine residues (2.62 x 10^5^) in the human proteome. Note, these numbers are different from the unique values because they include repeats: multiple copies of specific oxiforms.

In sum, as a rule tantamount to law: *N_c_* = *N_i_*. While *Ni* will be displaced from thermodynamic equilibrium towards the reduced state (e.g., ≈90%-reduced), it is possible that the number of partially oxidised cysteine redox proteoform molecules is in the order of 10^6-8^ per HeLa cell.

### 2.6. Quantifying the accessed *i* space in biological systems

By theoretically estimating ≈10^6-8^ partially oxidised cysteine redox proteoforms per HeLa cell, we have neared the limit of what can be deduced about the biologically accessed *i* space from theory alone. To move beyond theory by gaining empirical insights on the biological accessed *i* space, we analysed studies that used the maleimide-conjugated polyethylene glycol (PEG) assay (m-PEG assay) [44–47]. The PEG-payloads mobility-shift oxiforms by their cysteine oxidation integer, generating a ladder of increasingly oxidised bands [48]. Many unique proteoforms can coinhabit the partially oxidised oxiform bands. For example, 252 unique proteoforms may coinhabit a 50%-oxidised band for a protein with 10 cysteine residues. For a protein with 10 cysteine residues, the m-PEG assay collapses 1,024 unique proteoforms into 11 bands, simplifying the *i* space by 2-logs. If 3 bands were observed, one can state that 27.3% of the percentage redox graded *i* space (%-*i*-space) was accessed. Excluding the 100%-reduced, one can state that 20% of the %-oxiform space was accessed.

Here, we analysed 19 m-PEG assay studies (supplementary data file 9). The studies provided data on 33 cases, spanning 30 unique proteins from 6 species including 17 human proteins. Our secondary analysis revealed that 65.1% (SD = 34.4%, range = 9.5-100%) of the theoretical %-*i*-space was biologically accessed (Figure 6A). As expected, every protein formed the 100%-reduced cysteine redox proteoform. Unexpectedly, the number of bands containing at least one oxidised cysteine residue was appreciable. Specifically, 60.7% (SD = 38.1%, range = 5-100%) of the theoretical %-oxiform bands were observed. In 20 cases (60.6%), the 100%-oxidised proteoform was formed. Their formation pointed to a rich cysteine redox proteoform landscape.

**Figure 6.**
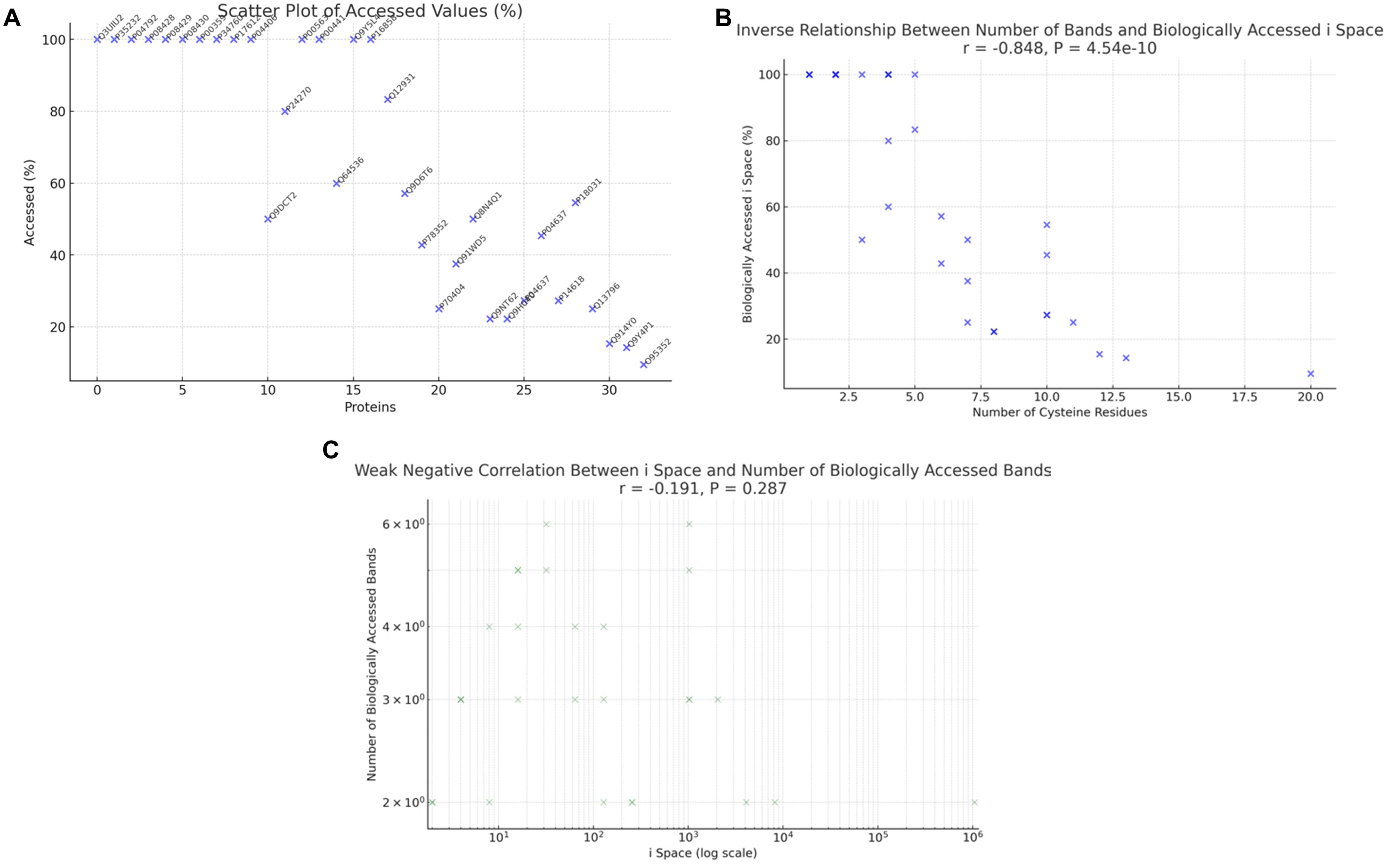
Quantifying the number of biologically accessed cysteine redox proteoforms using empirical data. A. Scatter Plot of Accessed Values (%): This plot shows the percentage of biologically accessed *i* space for each protein, identified by their UniProt IDs. B. Inverse Relationship Between Number of Bands and Biologically Accessed *i* Space: This plot depicts the strong inverse relationship between the number of cysteine residues (bands) and the biologically accessed *i* space percentage. The correlation coefficient (*r*) and p-value indicate a significant negative correlation. C. Weak Negative Correlation Between *i* Space and Number of Biologically Accessed Bands: This plot shows the weak negative correlation between the *i* space (on a log scale) and the number of biologically accessed bands. The correlation coefficient (*r*) and p-value indicate the relationship’s strength and significance.

Our analysis revealed two interesting features:

1. A strong inverse relationship (*r* = -0.848, *P* = 4.54 x 10^−10^) between the number of bands and the biologically accessed *i* space (Figure 6B). As the number of possible bands increased as a function of the cysteine residue integer, the percentage biologically accessed *i* space decreased. For example, 100% of the possible band space was sampled for every protein with 1-2 cysteine residues whereas just 5% was sampled for the [ATG7](https://www.uniprot.org/uniprotkb/O95352/entry) with 20 cysteines. This result is consistent with the equation 5 limiting the biologically accessible *i* space by 2-logs, from 1048576 to 13993 (-98.6%) unique ATG7-specific cysteine redox proteoforms, in a HeLa cell.
2. A weak negative correlation (*r* = -0.191, *P* = 0.287) between the *i* space and the number of biologically accessed bands (Figure 6C). Their relationship is weak because it is mathematically possible for a limited number of unique cysteine redox proteoforms to access all of the percentage redox graded *i* space. For example, 11 of 1,024 cysteine redox proteoforms are needed to access all of the percentage redox graded *i* space available to a protein with 10 cysteine residues. A minimum of 109 unique cysteine redox proteoforms can access over 65% of percentage redox grades available to 1.06 x 10^6^ cysteine redox proteoforms. Just 229 (0.02%) unique cysteine redox proteoforms would be needed to access the entire percentage redox graded space.

In sum, empirical data is broadly consistent with biological systems comprising a significant number of partially oxidised cysteine redox proteoforms. However, our conclusions are tempered by the paucity of empirical data. Meaning 99.92% of the human proteome is unmeasured. To allow their measurement formidable technical obstacles must be addressed [49–54]. For example, the m-PEG assay is rate-limited by the bulky PEG-payloads sterically blocking antibody binding [55–57]. We call for the concerted development of novel technologies for quantifying the biologically accessed *i* space.

### 2.7. Visualising proteoform-specified redox codes

When new technologies become available, exploring the biologically accessed *i* space may unmask a proteoform-specified code, such as cysteine oxidation hierarchies. To visualise these proteoform-specified codes, we created a streamlit app for generating <u>c</u>ysteine redox <u>p</u>roteoform <u>d</u>iagrams (CPDs) for a given protein using a Uniprot accession ([CPD_generator](https://cpdgeneratorispace.streamlit.app/)). To spotlight a CPD, we plotted the theoretical %-*i*-space in a histogram and immunoblot format for human [ATP5A](https://www.uniprot.org/uniprotkb/P25705/entry) as a redox regulated protein [58] amenable to m-PEG analysis. Since every band is sampled in *X. laevis* oocytes [47,59,60], we defined the potential paths to the ATP5A-specific proteoform bands for the two conserved cysteines Cys244 (C1) and Cys294 (C2) in the CPD (Figure 7).

**Figure 7.**
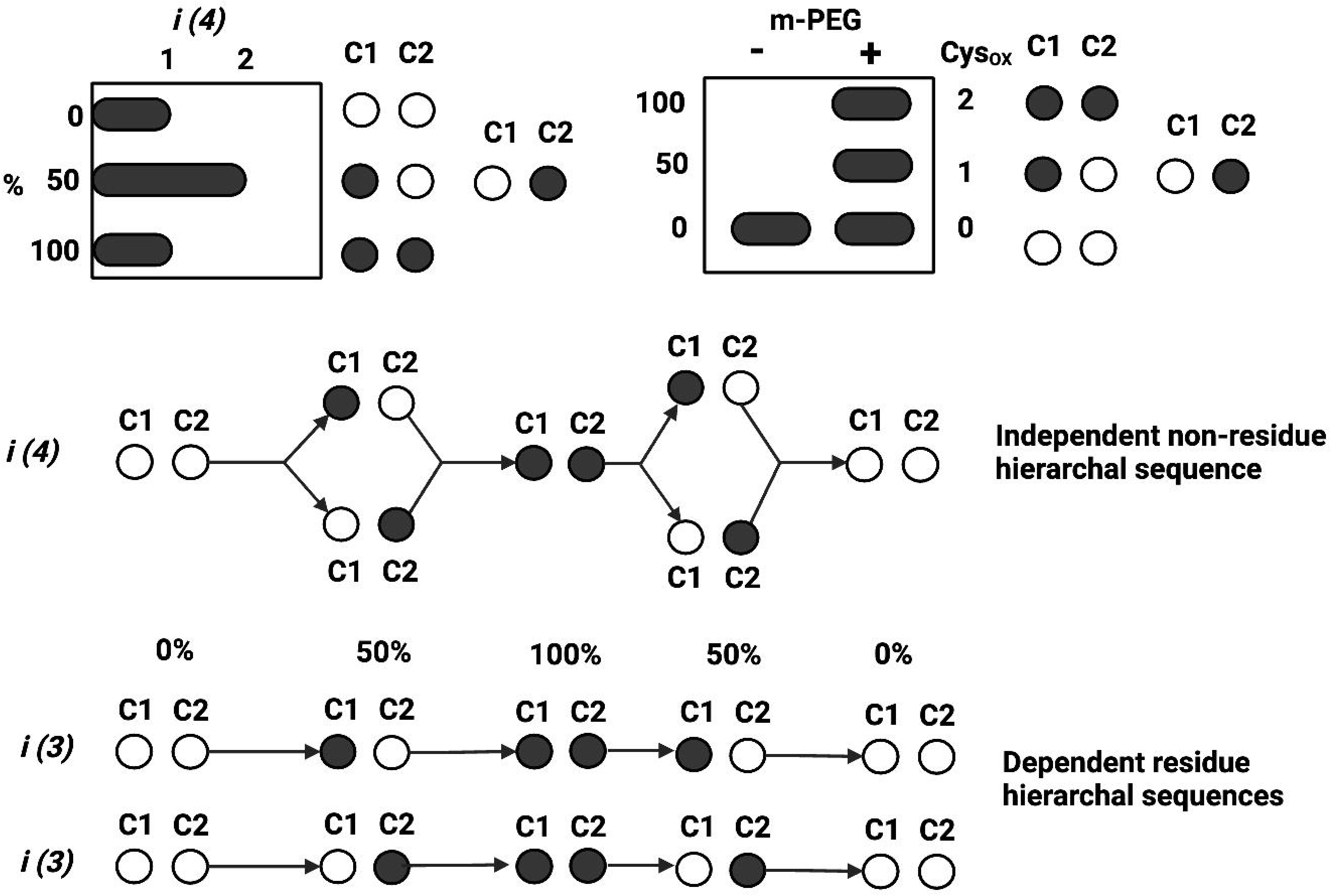
The top left plot depicts the distribution of unique cysteine redox proteoforms (*i*) for a protein with 2 cysteine residues (*n^R^* = 4), such as ATP5A, across the three mathematically allowed percentage redox grades. The white and black circles represent reduced and oxidised cysteines residues (C1: Cys244 or C2: Cys294), respectively. The top right plot depicts an outcome of a maleimide-conjugated polyethylene glycol (PEG)-payload (m-PEG) immunoblot assay. Without m-PEG, all 4 proteoforms migrate at an identical apparent molecular mass. With m-PEG, each oxiform, a proteoform containing at least one oxidised cysteine, is electrically mobility-shifted to an extent that depends on the number of oxidised cysteine residues. The 100%-reduced and 100%-oxidised binary bands are cysteine residue specified. The lower panel displays <u>c</u>ysteine redox <u>p</u>roteoform <u>d</u>iagrams (CPD). They illustrate the cysteine residue independent and dependent pathways in binary redox phase space. In the independent pathway (top CPD), the oxidation of C1 is not conditional upon the oxidation of C2 and vice versa. With no hierarchy between the residues, all the biologically accessible paths between the different percentage redox grades can be explored. In dependent pathways, the stepwise movement between percentage redox grades is conditional upon the oxidation of C1 (middle CPD) or C2 (lower CPD). By introducing hierarchal dependence into the sequence conditional rules generate preferential oxidation pathways that minimise the accessed *i* space.

In the independent pathway (top CPD), C1 oxidation is not reliant C2 oxidation and vice versa. Independence means all of the possible paths between the distinct %-*i*-space can be explored, maximising the biologically accessed *i* space. The implications for a redox biological function (*f*) in binary reduced (0) and oxidised (1) terms are:

*f* = C1_1_/C2_0_ OR C1_0_/C2_1_ where *f* depends on just the 50% redox grade (i.e., *f* is cysteine residue agnostic).

*f* = C1_1_ & C2_1_ via either 50% redox grade pathway where *f* depends on the 100% redox grade (i.e., there is no preferred route to *f*).

In the dependent pathway, stepwise movements in %-*i*-space depend upon the oxidation of C1 (middle CPD) or C2 (lower CPD). Hierarchal cysteine oxidation dependencies generate preferred pathways that minimise the biologically accessed *i* space. The implications for *f* in redox binary are:

*f* = C1_1_/C2_0_ where *f* depends on just the 50% redox grade (i.e., *f* is cysteine residue dependent).

*f* = C1_1_ & C2_1_ via C1_1_/C2_0_ 50%-redox grade pathway where *f* depends on the 100% redox grade (i.e., *f* is route dependent).

Equally, stepwise reductive movements in %-*i*-space can be dependent or independent. Different rules may operate in the “oxidative” and “reductive” branches of the CPD path space. For example, the forward “oxidative direction” path to 100%-oxidised proteoform may rely on C1 priming C2 oxidation but the backward “reductive direction” path to the 100%-reduced proteoform may be independent. The “start” of the path may be 50%-oxidised, with cyclical up or down redox steps from there. This point introduces the idea of setpoints and multiples thereof for different pools of the same molecule that are encoded by the biological context, such as proximity to an enzyme like [thioredoxin 1](https://www.uniprot.org/uniprotkb/P10599/entry). Like the strings of an instrument, setpoints may constrain the “notes” that otherwise identical molecules play, affording more or less access to a proteoform-based redox “key” range.

Numerous, nuanced proteoform-level redox codes, with all their intricacies, are possible with higher cysteine residue counts. In the independent pathway, the number of permutations specifying a 100%-oxidised *f* equals *i*. For example, 1,024 different paths to the 100%-oxidised state are possible for a protein with 10 cysteine residues. Defined dependencies can cap the path length for proteins with higher cysteine residue counts. For example, a strict conditional linear sequence would constrain the biologically accessed *i* space to 11 unique cysteine redox proteoforms (i.e., 10 processive redox steps along the path). A value between 11-1,024 may be traversed depending on the nature of the redox code. In sum, CPDs visualise proteoform-specified redox codes.

## 3. Conclusion

As Table 4 attests, the present study advanced current knowledge of cysteine redox proteoforms [1]. For example, the heterogenous distribution of the biologically accessible *i* space suggests some fundamental design principles, such as a constricted *i* space for abundant proteins favouring the formation of the desired oxiforms. By providing reference values, equations, and data files, the present work is expected to facilitate the exploration of the unmapped cysteine redox proteoform landscape. Ultimately, exploring the landscape of proteoforms that do form is one of the most important tasks in modern redox biology for these proteoforms actuate cysteine-dependent protein function, redox regulation, and oxidative stress.

**Table 4.**
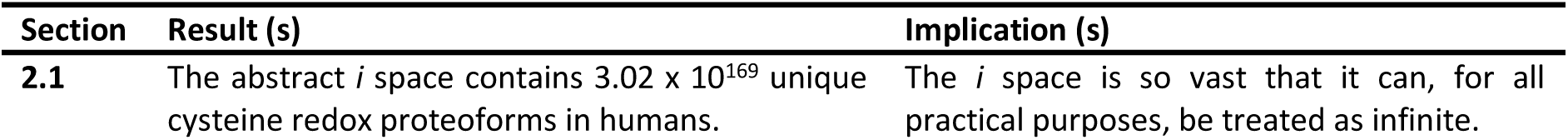
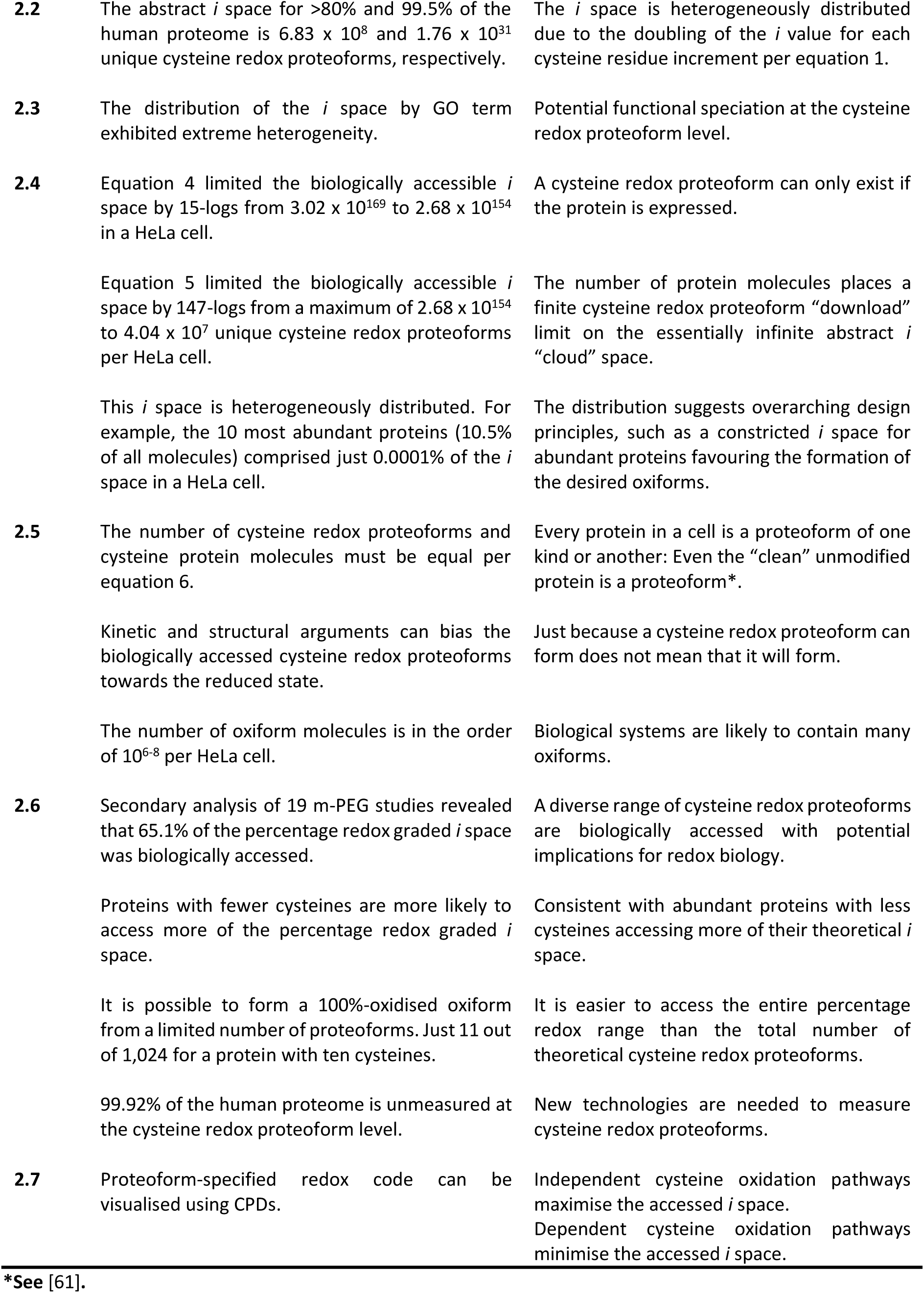
Summary of the main results of the present work and their implications by section.

## 4. Methods

### 4.1. Coding environment and repository

All computational analyses were performed using [Python](https://www.python.org/) in a Google Colaboratory ([Colab](https://colab.google/)) environment, which provided an interactive workspace for executing Python code. The main libraries used include BioPython, Pandas, Numpy, and Scipy, among others (requirements.txt). The source code is available in our GitHub repository https://github.com/JamesCobley/Cysteine-i-cloud inclusive of a [readme](https://github.com/JamesCobley/Cysteine-i-cloud/blob/main/README.md) file. The source code is freely available under a MIT license, allowing for broad use and modification. The source code can be cited using the Zendo DOI.

### 4.2. Calculating the theoretical *i* space of reference proteomes

To calculate the theoretical *i* space, reference proteomes (Table 5) were downloaded from Uniprot as gzipped FATSA files, which were uploaded onto a Colab workspace where the scriptcys1.py code was implemented. The script pre-processed the files by reading the gzipped FASTA file and extracting the amino acid sequences. Header lines, which started with the ’>’ character, were skipped to ensure only the sequence data were processed. Each line was stripped of trailing newlines or spaces. After importing the BioPython and pandas libraries, the script:

1. Parsed the gzip file and extract protein sequences.
2. Counted the number of cysteine residues (’C’) in each protein sequence.
3. Listed the number of proteins with different counts of cysteine residues.
4. Exported the results to an Excel file with two columns: column A for cysteine residues and column B for the number of proteins.

**Table 5.** Downloaded reference proteomes by species with strain information where applicable.

| Species | Reference proteome | Strain |
| --- | --- | --- |
| *E. coli* | https://www.uniprot.org/proteomes?query=%28organism_id%3A83333%29+AND+%28proteome_type%3A1%29 | Strain K12 (83333) |
| *S. cerevisiae* | https://www.uniprot.org/uniprotkb?query=yeast&facets=reviewed%3Atrue%2Cmodel_organism%3A559292 | strain ATCC 204508 / S288c |
| *C. elegans* | https://www.uniprot.org/proteomes/UP000001940 | Bristol N2 |
| *Drosophila* | https://www.uniprot.org/proteomes/UP000000803 | Berkeley |
| *D. rerio* | https://www.uniprot.org/proteomes/UP000000437 | n/a |
| *X. tropicalis* | https://www.uniprot.org/proteomes/UP000008143 | n/a |
| *M. musculus* | https://www.uniprot.org/proteomes/UP000000589 | C57BL/6J |
| *H. sapiens* | https://www.uniprot.org/uniprotkb?query=human&facets=model_organism%3A9606%2Creviewed%3Atrue | n/a |

The Excel file was processed by implementing a “PERMUTATONIA” function to solve equation 1 as “*i space*” for each “*cysteine residue*” value. For example, 2 = 4. Equation 3 was implemented by multiplying “*i space*” by “*proteins*” (e.g., 4 x 10 = 40). The resultant values were before summed to calculate the theoretical *i* space.

### 4.3. Defining the composition of the human *i* space

To define the composition of the human *i* space, the scriptcys1.py script output file (supplementary data file 1) was used to create the visualisations displayed in Figure 2 (section 4.9). To retrieve the identity of the protein with the most cysteine residues, we used the Cys_R_integer_find.py script. Supplementary data file 1 were manually analysed to create Table 2 by sorting the data using cysteine classifier ranges.

### 4.4. Defining the distribution of the human *i* space using GO terms

To define the distribution of human *i* space using GO terms, the custom Cys-count-ID.py script was implemented to categorise human proteins (identified by their UniProt accessions) by their cysteine residue counts. The output file, Supplementary Data File 3, organised the data such that each row corresponds to a UniProt accession, and each column represents a specific cysteine residue count. Generating the supplementary data file 3, enabled us to implement the Cys_GO_term.py script for sorting the human *i* space by selected GO terms. The *i* space for each term was computed by manually implementing equation 1 and 3 in excel per section 4.2.

### 4.5. Mathematically limiting the biological accessible *i* space

To mathematically limit the biologically accessible *i* space, equation 4 constrained the number of “downloadable” cysteine redox proteoforms using a protein expression argument. To illustrate equation 4, the custom Cys_Expression.py script was implemented on empirical data (supplementary Table S2 from [32]). Equation 5 was implemented by calculating the *i* value for each protein per equation 1 and summing them to derive a biologically accessible *i* space value.

After designing it to limit the biologically accessible *i* space using a copy number (N) argument, we equation 5 on empirical data (supplementary Table S7 from [32]). The Cys_Expression.py was implemented to assign a cysteine residue integer value to each N value before an *i* value was derived. An Excel “MIN” function was used to select the minimum value of *N* or *i*. The minimum values were then summed to compute the biologically accessible *i* space per equation 5.

### 4.6. Factors controlling the number of cysteine redox proteoforms that are biologically accessed

To illustrate a structural argument that could limit the biologically accessible *i* space, an Alphafold structure of human [PTEN](https://alphafold.ebi.ac.uk/entry/P60484) was uploaded into Colab. The custom Cys_SASA.py script was implemented to calculate and then print the SASA value for the sulfur atom (SG) and alpha carbon (CA) of the cysteine residues in PTEN.

To illustrate a kinetic argument that could limit the biologically accessible *i* space, we used PTEN. We first calculated the molarity of PTEN inside a HeLa cell by converting the *N* value in supplementary data file 8 to moles. To do so, an *N* of 3188 was divided 6.022 x 10^23^ and the HeLa cell volume of 2.6×10^3^ μm^3^ [62], equivalent to 2.6×10^−12^ L. This gave a molar value for PTEN of ≈2.04 nM. The oxidation of PTEN at Cys124 to give the RSOH was modelled as a bimolecular reaction between PTEN and H_2_O_2_ using an estimated *k* of 5.717 M^-1^ s^-1^ per equation 8.

**Equation 8.** *Rate* = *k x* [*PTEN*] *x* [*H*_2_*O*_2_]

The H_2_O_2_ concentration was assumed to be 10 nM. Substituting the known values for the PTEN concentration and the H_2_O_2_ concentration into the rate equation, the rate of Cys124 oxidation was calculated as:

**Equation 9.** 5.717 *x* 10^9^ *M*^−1^*s*^−1^ 2.04 *x* 10^−9^ *M*^−1^*s*^−1^ *x* 10 *x* 10 10^−9^ *M*^−1^*s*^−1^ = 1.166 *x* 10^−7^ *M*^−1^*s*^−1^

To estimate the number of oxidised PTEN molecules over 1 h (3600 s), we used the first-order reaction equation:

**Equation 10.** *Fraction oxidised* = 1 − *e*^*kt*^ where k =1.166×10^−7^ s^−1^ and t = 3600 s

The fraction of Cys124-oxidised PTEN molecules was calculated to be approximately 0.0004195. Multiplying this fraction by the total number of PTEN molecules in the cell (3188 molecules), the expected number of oxidised PTEN molecules was determined to be approximately 1.34 molecules (1 molecule). To exploit precise floating-point arithmetic, these calculations were also automated using the custom Cys_k_est.py script.

### 4.7. Quantifying the accessed *i* space in biological systems

To quantify the accessed *i* space in biological systems using empirical m-PEG assay data [59], relevant papers from the [PubMed](https://pubmed.ncbi.nlm.nih.gov/) database were manually retrieved using search terms, such as “maleimide-PEG”. Since the m-PEG assay was introduced in 2001 [44], the database was searched from 2001-2024. To maximise the number of papers retrieved, papers citing selected papers were manually parsed on [Google Scholar](https://scholar.google.com/). Given the focus on humans, papers measuring plant proteins were excluded. After retrieving the papers, each one was manually inspected to extract the data in Supplementary Data File 9, such as the number of bands that were biologically accessed.

### 4.8. Visualising proteoform-specified codes

To visualise proteoform-specified codes, we created the custom Cys_CPD_code.py script, located in a distinct repository https://github.com/JamesCobley/CPD, to run on the streamlit [application](https://cpdgeneratorispace.streamlit.app/). To use the application, one inputs a Uniprot accession for a protein of interest. The script prints the number of cysteine residues, their positions, and the number of cysteine redox proteoforms. In addition, the script prints the number of cysteine redox proteoforms in each percentage redox grade using Binomial theorem. Finally, the script generates a heatmap visualisation, a CPD, of the cysteine redox proteoforms. Reduced and oxidised cysteines are displayed in white and black, respectively.

### 4.9. Visualisations

The plots displayed to visualise the data were created in colab using the Matlab package and exported at 300 DPI.

### 4.10. Summary statistics and statistical analysis

Certain summary values were calculated using automated scripts. For example, the number of cysteines in the human proteome was calculated using the script summaryscript.py. After importing numpy into colab, most summary statistics, such as mean and median values, were calculated using scripts, such as [Cysteinestats.py](https://github.com/JamesCobley/Cysteine-i-cloud/blob/main/summaryscript.py). In other cases, summary statistics, such as the standard deviation, were calculated using excel functions. Correlation analyses were performed using the scipy package with alpha <0.05.

## Acknowledgements

Figure 7 was created using [Biorender](https://www.biorender.com/) and exported with a publication permission license. We thank Professor Angus Lamond (Dundee University) and all of the members of the Lamond lab for helpful scientific discussions.

## Author contributions

**James N. Cobley:** Conceptualisation, Methodology, Software; Data curation, Investigation, Visualisation, Writing -Original Draft Writing-review and editing. **Panagiotis N. Chatzinikolaou:** Resources, Software, Visualisation, Writing-review and editing. **Cameron Schmidt:** Resources, Software, Visualisation, Writing-review and editing.

## References

[1] J.N. Cobley, Oxiforms: Unique cysteine residue-and chemotype-specified chemical combinations can produce functionally-distinct proteoforms, Bioessays 45 (2023). 10.1002/bies.202200248.

[2] L.M. Smith, N.L. Kelleher, M. Linial, D. Goodlett, P. Langridge-Smith, Y.A. Goo, G. Safford, L. Bonilla*, G. Kruppa, R. Zubarev, J. Rontree, J. Chamot-Rooke, J. Garavelli, A. Heck, J. Loo, D. Penque, M. Hornshaw, C. Hendrickson, L. Pasa-Tolic, C. Borchers, D. Chan, N. Young*, J. Agar, C. Masselon, M. Gross*, F. McLafferty, Y. Tsybin, Y. Ge, I. Sanders*, J. Langridge, J. Whitelegge*, A. Marshall, Proteoform: a single term describing protein complexity, Nat Methods 10 (2013) 186–187. 10.1038/nmeth.2369.

[3] L.M. Smith, N.L. Kelleher, Proteoforms as the next proteomics currency, Science 359 (2018) 1106–1107. 10.1126/science.aat1884.

[4] R. Aebersold, J.N. Agar, I.J. Amster, M.S. Baker, C.R. Bertozzi, E.S. Boja, C.E. Costello, B.F. Cravatt, C. Fenselau, B.A. Garcia, Y. Ge, J. Gunawardena, R.C. Hendrickson, P.J. Hergenrother, C.G. Huber, A.R. Ivanov, O.N. Jensen, M.C. Jewett, N.L. Kelleher, L.L. Kiessling, N.J. Krogan, M.R. Larsen, J.A. Loo, R.R.O. Loo, E. Lundberg, M.J. MacCoss, P. Mallick, V.K. Mootha, M. Mrksich, T.W. Muir, S.M. Patrie, J.J. Pesavento, S.J. Pitteri, H. Rodriguez, A. Saghatelian, W. Sandoval, H. Schlüter, S. Sechi, S.A. Slavoff, L.M. Smith, M.P. Snyder, P.M. Thomas, M. Uhlén, J.E.V. Eyk, M. Vidal, D.R. Walt, F.M. White, E.R. Williams, T. Wohlschlager, V.H. Wysocki, N.A. Yates, N.L. Young, B. Zhang, How many human proteoforms are there?, Nat Chem Biol 14 (2018) 206–214. 10.1038/nchembio.2576.

[5] Y.-M. Go, J.D. Chandler, D.P. Jones, The cysteine proteome, Free Radical Bio Med 84 (2015) 227–245. 10.1016/j.freeradbiomed.2015.03.022.

[6] C.E. Paulsen, K.S. Carroll, Cysteine-Mediated Redox Signaling: Chemistry, Biology, and Tools for Discovery, Chem Rev 113 (2013) 4633– 4679. 10.1021/cr300163e.

[7] S. Parvez, M.J.C. Long, J.R. Poganik, Y. Aye, Redox Signaling by Reactive Electrophiles and Oxidants, Chem Rev 118 (2018) 8798–8888. 10.1021/acs.chemrev.7b00698.

[8] B. D’Autréaux, M.B. Toledano, ROS as signalling molecules: mechanisms that generate specificity in ROS homeostasis, Nat Rev Mol Cell Bio 8 (2007) 813–824. 10.1038/nrm2256.

[9] K.M. Holmström, T. Finkel, Cellular mechanisms and physiological consequences of redox-dependent signalling, Nat Rev Mol Cell Bio 15 (2014) 411–421. 10.1038/nrm3801.

[10] H. Sies, D.P. Jones, Reactive oxygen species (ROS) as pleiotropic physiological signalling agents, Nat Rev Mol Cell Bio 21 (2020) 363–383. 10.1038/s41580-020-0230-3.

[11] H. Sies, R.J. Mailloux, U. Jakob, Fundamentals of redox regulation in biology, Nat. Rev. Mol. Cell Biol. (2024) 1–19. 10.1038/s41580-024-00730-2.

[12] C. Lennicke, H.M. Cochemé, Redox metabolism: ROS as specific molecular regulators of cell signaling and function, Mol Cell 81 (2021) 3691–3707. 10.1016/j.molcel.2021.08.018.

[13] R.L.M. van Montfort, M. Congreve, D. Tisi, R. Carr, H. Jhoti, Oxidation state of the active-site cysteine in protein tyrosine phosphatase 1B, Nature 423 (2003) 773–777. 10.1038/nature01681.

[14] A. Salmeen, J.N. Andersen, M.P. Myers, T.-C. Meng, J.A. Hinks, N.K. Tonks, D. Barford, Redox regulation of protein tyrosine phosphatase 1B involves a sulphenyl-amide intermediate, Nature 423 (2003) 769–773. 10.1038/nature01680.

[15] B. Halliwell, J. Gutteridge, Free Radicals in Biology and Medicine, 2015.

[16] B.C. Dickinson, C.J. Chang, Chemistry and biology of reactive oxygen species in signaling or stress responses, Nat Chem Biol 7 (2011) 504–511. 10.1038/nchembio.607.

[17] C.C. Winterbourn, Reconciling the chemistry and biology of reactive oxygen species, Nat Chem Biol 4 (2008) 278–286. 10.1038/nchembio.85.

[18] H. Sies, C. Berndt, D.P. Jones, Oxidative Stress, Annu Rev Biochem 86 (2016) 1–34. 10.1146/annurev-biochem-061516-045037.

[19] H. Sies, Oxidative Eustress: On Constant Alert for Redox Homeostasis, Redox Biol 41 (2021) 101867. 10.1016/j.redox.2021.101867.

[20] H. Sies, Oxidative stress: a concept in redox biology and medicine, Redox Biol 4 (2015) 180–183. 10.1016/j.redox.2015.01.002.

[21] J.N. Cobley, 50 shades of oxidative stress: A state-specific cysteine redox pattern hypothesis, Redox Biol. 67 (2023) 102936. 10.1016/j.redox.2023.102936.

[22] J.N. Cobley, Exploring the unmapped cysteine redox proteoform landscape, Am. J. Physiol.-Cell Physiol. (2024). 10.1152/ajpcell.00152.2024.

[23] I.H.G.S. Consortium, W.I. for B.R. Research: Center for Genome, E.S. Lander, L.M. Linton, B. Birren, C. Nusbaum, M.C. Zody, J. Baldwin, K. Devon, K. Dewar, M. Doyle, W. FitzHugh, R. Funke, D. Gage, K. Harris, A. Heaford, J. Howland, L. Kann, J. Lehoczky, R. LeVine, P. McEwan, K. McKernan, J. Meldrim, J.P. Mesirov, C. Miranda, W. Morris, J. Naylor, C. Raymond, M. Rosetti, R. Santos, A. Sheridan, C. Sougnez, N. Stange-Thomann, N. Stojanovic, A. Subramanian, D. Wyman, T.S. Centre:, J. Rogers, J. Sulston, R. Ainscough, S. Beck, D. Bentley, J. Burton, C. Clee, N. Carter, A. Coulson, R. Deadman, P. Deloukas, A. Dunham, I. Dunham, R. Durbin, L. French, D. Grafham, S. Gregory, T. Hubbard, S. Humphray, A. Hunt, M. Jones, C. Lloyd, A. McMurray, L. Matthews, S. Mercer, S. Milne, J.C. Mullikin, A. Mungall, R. Plumb, M. Ross, R. Shownkeen, S. Sims, W.U.G.S. Center, R.H. Waterston, R.K. Wilson, L.W. Hillier, J.D. McPherson, M.A. Marra, E.R. Mardis, L.A. Fulton, A.T. Chinwalla, K.H. Pepin, W.R. Gish, S.L. Chissoe, M.C. Wendl, K.D. Delehaunty, T.L. Miner, A. Delehaunty, J.B. Kramer, L.L. Cook, R.S. Fulton, D.L. Johnson, P.J. Minx, S.W. Clifton, U.D.J.G. Institute:, T. Hawkins, E. Branscomb, P. Predki, P. Richardson, S. Wenning, T. Slezak, N. Doggett, J.-F. Cheng, A. Olsen, S. Lucas, C. Elkin, E. Uberbacher, M. Frazier, B.C. of M.H.G.S. Center:, R.A. Gibbs, D.M. Muzny, S.E. Scherer, J.B. Bouck, E.J. Sodergren, K.C. Worley, C.M. Rives, J.H. Gorrell, M.L. Metzker, S.L. Naylor, R.S. Kucherlapati, D.L. Nelson, G.M. Weinstock, R.G.S. Center:, Y. Sakaki, A. Fujiyama, M. Hattori, T. Yada, A. Toyoda, T. Itoh, C. Kawagoe, H. Watanabe, Y. Totoki, T. Taylor, G. and C. UMR-8030:, J. Weissenbach, R. Heilig, W. Saurin, F. Artiguenave, P. Brottier, T. Bruls, E. Pelletier, C. Robert, P. Wincker. D. of G.A. Biotechnology: Institute of Molecular, A. Rosenthal, M. Platzer, G. Nyakatura, S. Taudien, A. Rump, G.S. Center:, D.R. Smith, L. Doucette-Stamm, M. Rubenfield, K. Weinstock, H.M. Lee, J. Dubois, B.G.I.G. Center:, H. Yang, J. Yu, J. Wang, G. Huang, J. Gu, M.S.C. Biology: The Institute for Systems, L. Hood, L. Rowen, A. Madan, S. Qin, S.G.T. Center:, R.W. Davis, N.A. Federspiel, A.P. Abola, M.J. Proctor, U. of O.A.C. for G. Technology:, B.A. Roe, F. Chen, H. Pan, M.P.I. for M. Genetics:, J. Ramser, H. Lehrach, R. Reinhardt, C.S.H.L. Center: Lita Annenberg Hazen Genome, W.R. McCombie, M. de la Bastide, N. Dedhia, G.R.C. for Biotechnology:, H. Blöcker, K. Hornischer, G. Nordsiek, R. Agarwala, L. Aravind, J.A. Bailey, A. Bateman, S. Batzoglou, E. Birney, P. Bork, D.G. Brown, C.B. Burge, L. Cerutti, H.-C. Chen, D. Church, M. Clamp, R.R. Copley, T. Doerks, S.R. Eddy, E.E. Eichler, T.S. Furey, J. Galagan, J.G.R. Gilbert, C. Harmon, Y. Hayashizaki, D. Haussler, H. Hermjakob, K. Hokamp, W. Jang, L.S. Johnson, T.A. Jones, S. Kasif, A. Kaspryzk, S. Kennedy, W.J. Kent, P. Kitts, E.V. Koonin, I. Korf, D. Kulp, D. Lancet, T.M. Lowe, A. McLysaght, T. Mikkelsen, J.V. Moran, N. Mulder, V.J. Pollara, C.P. Ponting, G. Schuler, J. Schultz, G. Slater, A.F.A. Smit, E. Stupka, J. Szustakowki, D. Thierry-Mieg, J. Thierry-Mieg, L. Wagner, J. Wallis, R. Wheeler, A. Williams, Y.I. Wolf, K.H. Wolfe, S.-P. Yang, R.-F. Yeh, S. management: N.H.G.R.I. Health: US National Institutes of, F. Collins, M.S. Guyer, J. Peterson, A. Felsenfeld, K.A. Wetterstrand, S.H.G. Center:, R.M. Myers, J. Schmutz, M. Dickson, J. Grimwood, D.R. Cox, U. of W.G. Center:, M.V. Olson, R. Kaul, C. Raymond, D. of M.B. Medicine: Keio University School of, N. Shimizu, K. Kawasaki, S. Minoshima, U. of T.S.M.C. at Dallas:, G.A. Evans, M. Athanasiou, R. Schultz, O. of S. Energy: US Department of, A. Patrinos, T.W. Trust:, M.J. Morgan, Initial sequencing and analysis of the human genome, Nature 409 (2001) 860–921. 10.1038/35057062.

[24] J.C. Venter, M.D. Adams, E.W. Myers, P.W. Li, R.J. Mural, G.G. Sutton, H.O. Smith, M. Yandell, C.A. Evans, R.A. Holt, J.D. Gocayne, P. Amanatides, R.M. Ballew, D.H. Huson, J.R. Wortman, Q. Zhang, C.D. Kodira, X.H. Zheng, L. Chen, M. Skupski, G. Subramanian, P.D. Thomas, J. Zhang, G.L.G. Miklos, C. Nelson, S. Broder, A.G. Clark, J. Nadeau, V.A. McKusick, N. Zinder, A.J. Levine, R.J. Roberts, M. Simon, C. Slayman, M. Hunkapiller, R. Bolanos, A. Delcher, I. Dew, D. Fasulo, M. Flanigan, L. Florea, A. Halpern, S. Hannenhalli, S. Kravitz, S. Levy, C. Mobarry, K. Reinert, K. Remington, J. Abu-Threideh, E. Beasley, K. Biddick, V. Bonazzi, R. Brandon, M. Cargill, I. Chandramouliswaran, R. Charlab, K. Chaturvedi, Z. Deng, V.D. Francesco, P. Dunn, K. Eilbeck, C. Evangelista, A.E. Gabrielian, W. Gan, W. Ge, F. Gong, Z. Gu, P. Guan, T.J. Heiman, M.E. Higgins, R.-R. Ji, Z. Ke, K.A. Ketchum, Z. Lai, Y. Lei, Z. Li, J. Li, Y. Liang, X. Lin, F. Lu, G.V. Merkulov, N. Milshina, H.M. Moore, A.K. Naik, V.A. Narayan, B. Neelam, D. Nusskern, D.B. Rusch, S. Salzberg, W. Shao, B. Shue, J. Sun, Z.Y. Wang, A. Wang, X. Wang, J. Wang, M.-H. Wei, R. Wides, C. Xiao, C. Yan, A. Yao, J. Ye, M. Zhan, W. Zhang, H. Zhang, Q. Zhao, L. Zheng, F. Zhong, W. Zhong, S.C. Zhu, S. Zhao, D. Gilbert, S. Baumhueter, G. Spier, C. Carter, A. Cravchik, T. Woodage, F. Ali, H. An, A. Awe, D. Baldwin, H. Baden, M. Barnstead, I. Barrow, K. Beeson, D. Busam, A. Carver, A. Center, M.L. Cheng, L. Curry, S. Danaher, L. Davenport, R. Desilets, S. Dietz, K. Dodson, L. Doup, S. Ferriera, N. Garg, A. Gluecksmann, B. Hart, J. Haynes, C. Haynes, C. Heiner, S. Hladun, D. Hostin, J. Houck, T. Howland, C. Ibegwam, J. Johnson, F. Kalush, L. Kline, S. Koduru, A. Love, F. Mann, D. May, S. McCawley, T. McIntosh, I. McMullen, M. Moy, L. Moy, B. Murphy, K. Nelson, C. Pfannkoch, E. Pratts, V. Puri, H. Qureshi, M. Reardon, R. Rodriguez, Y.-H. Rogers, D. Romblad, B. Ruhfel, R. Scott, C. Sitter, M. Smallwood, E. Stewart, R. Strong, E. Suh, R. Thomas, N.N. Tint, S. Tse, C. Vech, G. Wang, J. Wetter, S. Williams, M. Williams, S. Windsor, E. Winn-Deen, K. Wolfe, J. Zaveri, K. Zaveri, J.F. Abril, R. Guigó, M.J. Campbell, K.V. Sjolander, B. Karlak, A. Kejariwal, H. Mi, B. Lazareva, T. Hatton, A. Narechania, K. Diemer, A. Muruganujan, N. Guo, S. Sato, V. Bafna, S. Istrail, R. Lippert, R. Schwartz, B. Walenz, S. Yooseph, D. Allen, A. Basu, J. Baxendale, L. Blick, M. Caminha, J. Carnes-Stine, P. Caulk, Y.-H. Chiang, M. Coyne, C. Dahlke, A.D. Mays, M. Dombroski, M. Donnelly, D. Ely, S. Esparham, C. Fosler, H. Gire, S. Glanowski, K. Glasser, A. Glodek, M. Gorokhov, K. Graham, B. Gropman, M. Harris, J. Heil, S. Henderson, J. Hoover, D. Jennings, C. Jordan, J. Jordan, J. Kasha, L. Kagan, C. Kraft, A. Levitsky, M. Lewis, X. Liu, J. Lopez, D. Ma, W. Majoros, J. McDaniel, S. Murphy, M. Newman, T. Nguyen, N. Nguyen, M. Nodell, S. Pan, J. Peck, M. Peterson, W. Rowe, R. Sanders, J. Scott, M. Simpson, T. Smith, A. Sprague, T. Stockwell, R. Turner, E. Venter, M. Wang, M. Wen, D. Wu, M. Wu, A. Xia, A. Zandieh, X. Zhu, The Sequence of the Human Genome, Science 291 (2001) 1304–1351. 10.1126/science.1058040.

[25] M.-S. Kim, S.M. Pinto, D. Getnet, R.S. Nirujogi, S.S. Manda, R. Chaerkady, A.K. Madugundu, D.S. Kelkar, R. Isserlin, S. Jain, J.K. Thomas, B. Muthusamy, P. Leal-Rojas, P. Kumar, N.A. Sahasrabuddhe, L. Balakrishnan, J. Advani, B. George, S. Renuse, L.D.N. Selvan, A.H. Patil, V. Nanjappa, A. Radhakrishnan, S. Prasad, T. Subbannayya, R. Raju, M. Kumar, S.K. Sreenivasamurthy, A. Marimuthu, G.J. Sathe, S. Chavan, K.K. Datta, Y. Subbannayya, A. Sahu, S.D. Yelamanchi, S. Jayaram, P. Rajagopalan, J. Sharma, K.R. Murthy, N. Syed, R. Goel, A.A. Khan, S. Ahmad, G. Dey, K. Mudgal, A. Chatterjee, T.-C. Huang, J. Zhong, X. Wu, P.G. Shaw, D. Freed, M.S. Zahari, K.K. Mukherjee, S. Shankar, A. Mahadevan, H. Lam, C.J. Mitchell, S.K. Shankar, P. Satishchandra, J.T. Schroeder, R. Sirdeshmukh, A. Maitra, S.D. Leach, C.G. Drake, M.K. Halushka, T.S.K. Prasad, R.H. Hruban, C.L. Kerr, G.D. Bader, C.A. Iacobuzio-Donahue, H. Gowda, A. Pandey, A draft map of the human proteome, Nature 509 (2014) 575–581. 10.1038/nature13302.

[26] S. Adhikari, E.C. Nice, E.W. Deutsch, L. Lane, G.S. Omenn, S.R. Pennington, Y.-K. Paik, C.M. Overall, F.J. Corrales, I.M. Cristea, J.E.V. Eyk, M. Uhlén, C. Lindskog, D.W. Chan, A. Bairoch, J.C. Waddington, J.L. Justice, J. LaBaer, H. Rodriguez, F. He, M. Kostrzewa, P. Ping, R.L. Gundry, P. Stewart, S. Srivastava, S. Srivastava, F.C.S. Nogueira, G.B. Domont, Y. Vandenbrouck, M.P.Y. Lam, S. Wennersten, J.A. Vizcaino, M. Wilkins, J.M. Schwenk, E. Lundberg, N. Bandeira, G. Marko-Varga, S.T. Weintraub, C. Pineau, U. Kusebauch, R.L. Moritz, S.B. Ahn, M. Palmblad, M.P. Snyder, R. Aebersold, M.S. Baker, A high-stringency blueprint of the human proteome, Nat Commun 11 (2020) 5301. 10.1038/s41467-020-19045-9.

[27] V. Marx, Inside the chase after those elusive proteoforms, Nat. Methods (2024) 1–6. 10.1038/s41592-024-02170-4.

[28] I. Lorenzen, J.A. Eble, E.-M. Hanschmann, Thiol switches in membrane proteins -Extracellular redox regulation in cell biology, Biol Chem 402 (2021) 253–269. 10.1515/hsz-2020-0266.

[29] T. Yan, L.M. Boatner, L. Cui, P.J. Tontonoz, K.M. Backus, Defining the Cell Surface Cysteinome Using Two-Step Enrichment Proteomics, JACS Au 3 (2023) 3506–3523. 10.1021/jacsau.3c00707.

[30] S. Gobron, H. Monnerie, R. Meiniel, I. Creveaux, W. Lehmann, D. Lamalle, B. Dastugue, A. Meiniel, SCO-spondin: a new member of the thrombospondin family secreted by the subcommissural organ is a candidate in the modulation of neuronal aggregation, J. Cell Sci. 109 (1996) 1053–1061. 10.1242/jcs.109.5.1053.

[31] J.N. Cobley, M.L. Fiorello, D.M. Bailey, 13 reasons why the brain is susceptible to oxidative stress, Redox Biol 15 (2018) 490–503. 10.1016/j.redox.2018.01.008.

[32] N. Nagaraj, J.R. Wisniewski, T. Geiger, J. Cox, M. Kircher, J. Kelso, S. Pääbo, M. Mann, Deep proteome and transcriptome mapping of a human cancer cell line, Mol. Syst. Biol. 7 (2011) 548–548. 10.1038/msb.2011.81.

[33] H. Xiao, M.P. Jedrychowski, D.K. Schweppe, E.L. Huttlin, Q. Yu, D.E. Heppner, J. Li, J. Long, E.L. Mills, J. Szpyt, Z. He, G. Du, R. Garrity, A. Reddy, L.P. Vaites, J.A. Paulo, T. Zhang, N.S. Gray, S.P. Gygi, E.T. Chouchani, A Quantitative Tissue-Specific Landscape of Protein Redox Regulation during Aging, Cell 180 (2020) 968–983.e24. 10.1016/j.cell.2020.02.012.

[34] G.C. Brown, Bioenergetic myths of energy transduction in eukaryotic cells, Front. Mol. Biosci. 11 (2024) 1402910. 10.3389/fmolb.2024.1402910.

[35] D.P. Jones, H. Sies, The Redox Code, Antioxid Redox Sign 23 (2015) 734–746. 10.1089/ars.2015.6247.

[36] R.E. Hansen, D. Roth, J.R. Winther, Quantifying the global cellular thiol–disulfide status, Proc National Acad Sci 106 (2009) 422–427. 10.1073/pnas.0812149106.

[37] J. Jumper, R. Evans, A. Pritzel, T. Green, M. Figurnov, O. Ronneberger, K. Tunyasuvunakool, R. Bates, A. Žídek, A. Potapenko, A. Bridgland, C. Meyer, S.A.A. Kohl, A.J. Ballard, A. Cowie, B. Romera-Paredes, S. Nikolov, R. Jain, J. Adler, T. Back, S. Petersen, D. Reiman, E. Clancy, M. Zielinski, M. Steinegger, M. Pacholska, T. Berghammer, S. Bodenstein, D. Silver, O. Vinyals, A.W. Senior, K. Kavukcuoglu, P. Kohli, D. Hassabis, Highly accurate protein structure prediction with AlphaFold, Nature 596 (2021) 583–589. 10.1038/s41586-021-03819-2.

[38] T.C. Terwilliger, D. Liebschner, T.I. Croll, C.J. Williams, A.J. McCoy, B.K. Poon, P.V. Afonine, R.D. Oeffner, J.S. Richardson, R.J. Read, P.D. Adams, AlphaFold predictions are valuable hypotheses and accelerate but do not replace experimental structure determination, Nat. Methods 21 (2024) 110–116. 10.1038/s41592-023-02087-4.

[39] F. Antunes, P.M. Brito, Quantitative biology of hydrogen peroxide signaling, Redox Biol 13 (2017) 1–7. 10.1016/j.redox.2017.04.039.

[40] S.-R. Lee, K.-S. Yang, J. Kwon, C. Lee, W. Jeong, S.G. Rhee, Reversible Inactivation of the Tumor Suppressor PTEN by H2O2 *, J Biol Chem 277 (2002) 20336–20342. 10.1074/jbc.m111899200.

[41] H. Sies, Hydrogen peroxide as a central redox signaling molecule in physiological oxidative stress: Oxidative eustress, Redox Biol 11 (2017) 613–619. 10.1016/j.redox.2016.12.035.

[42] Y. Bodnar, C.H. Lillig, Cysteinyl and methionyl redox switches: Structural prerequisites and consequences, Redox Biol. 65 (2023) 102832. 10.1016/j.redox.2023.102832.

[43] J.B. Behring, S. van der Post, A.D. Mooradian, M.J. Egan, M.I. Zimmerman, J.L. Clements, G.R. Bowman, J.M. Held, Spatial and temporal alterations in protein structure by EGF regulate cryptic cysteine oxidation, Sci Signal 13 (2020) eaay7315. 10.1126/scisignal.aay7315.

[44] L. Makmura, M. Hamann, A. Areopagita, S. Furuta, A. Muoz, J. Momand, Development of a Sensitive Assay to Detect Reversibly Oxidized Protein Cysteine Sulfhydryl Groups, Antioxid Redox Sign 3 (2001) 1105–1118. 10.1089/152308601317203611.

[45] J.R. Burgoyne, O. Oviosu, P. Eaton, The PEG-switch assay: A fast semi-quantitative method to determine protein reversible cysteine oxidation, J Pharmacol Toxicol 68 (2013) 297–301. 10.1016/j.vascn.2013.07.001.

[46] L.A.G. van Leeuwen, E.C. Hinchy, M.P. Murphy, E.L. Robb, H.M. Cochemé, Click-PEGylation – A mobility shift approach to assess the redox state of cysteines in candidate proteins, Free Radical Bio Med 108 (2017) 374–382. 10.1016/j.freeradbiomed.2017.03.037.

[47] J.N. Cobley, A. Noble, E. Jimenez-Fernandez, M.-T.V. Moya, M. Guille, H. Husi, Catalyst-free Click PEGylation reveals substantial mitochondrial ATP synthase sub-unit alpha oxidation before and after fertilisation, Redox Biol 26 (2019) 101258. 10.1016/j.redox.2019.101258.

[48] J.N. Cobley, N.V. Margaritelis, P.N. Chatzinikolaou, M.G. Nikolaidis, G.W. Davison, Ten “Cheat Codes” for Measuring Oxidative Stress in Humans, Antioxidants 13 (2024) 877. 10.3390/antiox13070877.

[49] J.N. Cobley, G.K. Sakellariou, H. Husi, B. McDonagh, Proteomic strategies to unravel age-related redox signalling defects in skeletal muscle, Free Radical Bio Med 132 (2019) 24–32. 10.1016/j.freeradbiomed.2018.09.012.

[50] A. Po, C.E. Eyers, Top-Down Proteomics and the Challenges of True Proteoform Characterization, J. Proteome Res. 22 (2023) 3663–3675. 10.1021/acs.jproteome.3c00416.

[51] K.E. Burnum-Johnson, T.P. Conrads, R.R. Drake, A.E. Herr, R. Iyengar, R.T. Kelly, E. Lundberg, M.J. MacCoss, A. Naba, G.P. Nolan, P.A. Pevzner, K.D. Rodland, S. Sechi, N. Slavov, J.M. Spraggins, J.E.V. Eyk, M. Vidal, C. Vogel, D.R. Walt, N.L. Kelleher, New Views of Old Proteins: Clarifying the Enigmatic Proteome, Mol Cell Proteomics 21 (2022) 100254. 10.1016/j.mcpro.2022.100254.

[52] D.L. Plubell, L. Käll, B.-J. Webb-Robertson, L.M. Bramer, A. Ives, N.L. Kelleher, L.M. Smith, T.J. Montine, C.C. Wu, M.J. MacCoss, Putting Humpty Dumpty Back Together Again: What Does Protein Quantification Mean in Bottom-Up Proteomics?, J. Proteome Res. 21 (2022) 891–898. 10.1021/acs.jproteome.1c00894.

[53] N. Siuti, N.L. Kelleher, Decoding protein modifications using top-down mass spectrometry, Nat. Methods 4 (2007) 817–821. 10.1038/nmeth1097.

[54] D.J. Muggeridge, D.R. Crabtree, A. Tuncay, I.L. Megson, G. Davison, J.N. Cobley, Exercise decreases PP2A-specific reversible thiol oxidation in human erythrocytes: Implications for redox biomarkers, Free Radical Bio Med 182 (2022) 73–78. 10.1016/j.freeradbiomed.2022.02.019.

[55] A. Noble, M. Guille, J.N. Cobley, ALISA: A microplate assay to measure protein thiol redox state, Free Radical Bio Med 174 (2021) 272–280. 10.1016/j.freeradbiomed.2021.08.018.

[56] A. Tuncay, A. Noble, M. Guille, J.N. Cobley, RedoxiFluor: A microplate technique to quantify target-specific protein thiol redox state in relative percentage and molar terms, Free Radical Bio Med 181 (2022) 118–129. 10.1016/j.freeradbiomed.2022.01.023.

[57] J.N. Cobley, H. Husi, Immunological Techniques to Assess Protein Thiol Redox State: Opportunities, Challenges and Solutions, Antioxidants 9 (2020) 315. 10.3390/antiox9040315.

[58] S.-B. Wang, D.B. Foster, J. Rucker, B. O’Rourke, D.A. Kass, J.E.V. Eyk, Redox Regulation of Mitochondrial ATP Synthase, Circ Res 109 (2011) 750–757. 10.1161/circresaha.111.246124.

[59] J. Cobley, A. Noble, R. Bessell, M. Guille, H. Husi, Reversible Thiol Oxidation Inhibits the Mitochondrial ATP Synthase in Xenopus laevis Oocytes, Antioxidants 9 (2020) 215. 10.3390/antiox9030215.

[60] A. Tuncay, D.R. Crabtree, D.J. Muggeridge, H. Husi, J.N. Cobley, Performance benchmarking microplate-immunoassays for quantifying target-specific cysteine oxidation reveals their potential for understanding redox-regulation and oxidative stress, Free Radical Bio Med 204 (2023) 252–265. 10.1016/j.freeradbiomed.2023.05.006.

[61] K. Carbonara, M. Andonovski, J.R. Coorssen, Proteomes Are of Proteoforms: Embracing the Complexity, Proteomes 9 (2021) 38. 10.3390/proteomes9030038.

[62] R. Milo, What is the total number of protein molecules per cell volume? A call to rethink some published values, Bioessays 35 (2013) 1050–1055. 10.1002/bies.201300066.

